# Latitudinal clines in climate and sleep patterns shape disease outcomes in *Drosophila melanogaster* infected by *Metarhizium anisopliae*

**DOI:** 10.1101/2024.10.10.617675

**Authors:** Mintong Nan, Jonathan B. Wang, Michail Siokis, Raymond J. St. Leger

## Abstract

Major latitudinal clines have been observed in *Drosophila melanogaster*, a human commensal that originated in tropical Africa and has spread world-wide to colonize temperate habitats. However, despite the significant impact of pathogens on species distribution, the influence of geographical factors on disease susceptibility remains poorly understood. This study investigated the effects of latitudinal clines and biomes on disease resistance using the common fly pathogen *Metarhizium anisopliae* and 43 global *Drosophila melanogaster* populations. The results showed that disease resistance was correlated with latitudinal gradients of sleep duration, temperature and humidity. While fungal diversity at tropical latitudes may drive enhanced defenses, the most disease-resistant males were also the most susceptible to desiccation, indicating potential trade-offs between abiotic stress resistance, necessary for survival in temperate habitats, and disease resistance. The study also found that sex, mating status, and sleep interacted with abiotic stresses to impact disease resistance, with longer-sleeping males and virgin flies surviving infections longer, and extra daytime sleep post-infection being protective, especially in the most resistant fly lines. These findings promote the idea that sleep and defense against disease are intertwined traits related to organismal fitness and subject to joint clinal evolution.

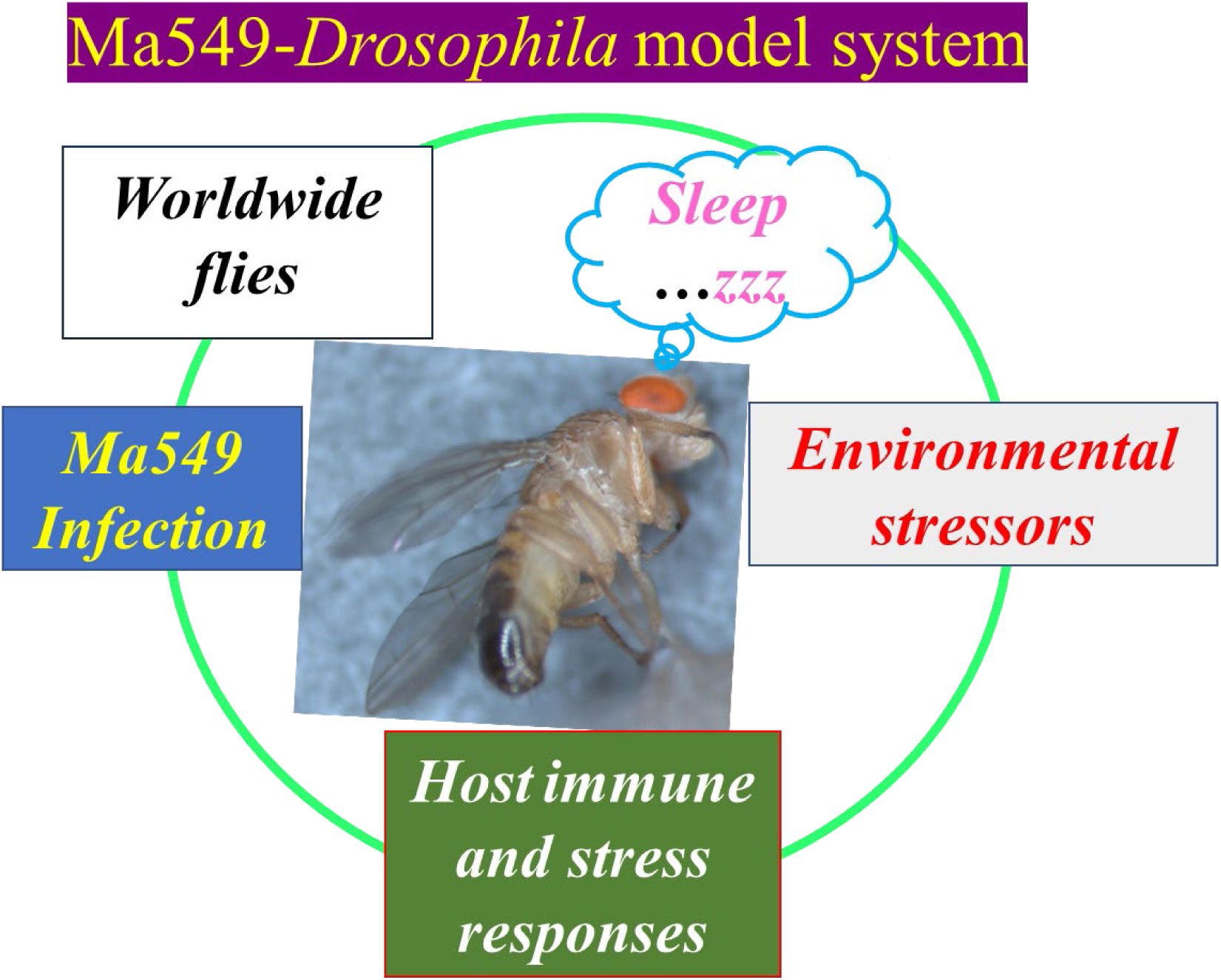

## Introduction

Understanding how organisms adapt to their environment, especially which biological processes are most relevant, is still an open question in biology (Mateo et al., 2018) and one that has become increasingly critical with growing threats to biodiversity from human activities. The fruit fly *Drosophila melanogaster*, a key evolutionary model, offers unique opportunities to investigate historical processes experienced by human commensals (Arguello et al., 2019). Originating in sub-Saharan Africa and initially specializing to marula fruit (*Sclerocarya birrea*), this species first associated with people when marula was stored by cave-dwellers (Mansourian et al., 2018). Flies maintained under laboratory conditions since 1916 still prefer marula over other fruits (Mansourian et al., 2018). *D. melanogaster* expanded its range to the Middle East around 13,000 years ago and to Europe and Asia about 1,800 years ago, coinciding with citrus and grape cultivation (Sprengelmeyer et al., 2019). Further colonization events, driven by human dispersal, have spread fly populations globally, including North America and Australia about 200 years ago (David & Capy, 1988; Haudry et al., 2020). Colonizing natural habitats, environmental clines, and geographic variables across continents suggests the capacity to rapidly adapt to local conditions (Blanford et al., 2003; Mateo et al., 2018), making *D. melanogaster* a robust model for studying evolutionary processes of range expansion and adaptation (Arguello et al., 2019) with promise for understanding evolutionary change in response to future environmental challenges. Thus, variations in *D. melanogaster* lines from diverse locales have identified sleep differences (Brown et al., 2018) linked to metabolic and foraging needs (Keene & Duboue) and associated with latitudinal clines and elevated temperatures (Brown et al., 2018). Other clines such as body size and chill resistance that vary predictably in abiotic factors have been the subject of much theoretical and experimental work but relatively little is known about clines whose evolution may be governed by interspecific interactions.

Flies potentially encounter distinct parasitic and pathogenic pressures in different environments, and according to the Red Queen hypothesis, hosts continually evolve to minimize the effects of pathogen adaptation (Blanford et al., 2003; Dybdahl & Lively, 1998; Lively & Dybdahl, 2000). Disease resistance varies across *Drosophila* species; those with a broad diet (e.g., *D. melanogaster, D. simulans, D. repleta, D. arizonae*) tend to be more resistant to *Metarhizium anisopliae* strain Ma549 than dietary specialists (e.g., *D. erecta, D. sechellia*) (O’Malley et al., 2023), possibly because insects with a broad diet have more opportunities to encounter and develop resistance to a variety of pathogens (O’Malley et al., 2023; Wang et al., 2017). There is also a great deal of intraspecific variation in disease resistance in several species (O’Malley et al., 2023; Wang et al., 2017), but whether any of this variation is predictable across environments is poorly understood. However, four African *D. melanogaster* lines were found to be more resistant to the fungal pathogen *Beauveria bassiana* than two non-African populations (Tinsley et al., 2006).

Overall, the ecological and evolutionary processes contributing to disease resistance are understudied. In contrast, there is a voluminous literature on the molecular mechanisms of host– pathogen interactions using *D. melanogaster* (Igboin et al., 2012; Lemaitre & Hoffmann, 2007) and these have shown a notable conservation of pathogenesis and host defense mechanisms between higher host organisms and *Drosophila* (Panayidou et al., 2014; Ugur et al., 2016). Nonetheless, both individual fruit flies and humans exhibit extensive variability in their responses to infectious agents (Lu & St Leger, 2016; Råberg et al., 2007), as demonstrated by human variability to COVID-19. Understanding why two individuals exposed to identical infections have different outcomes is a central issue in infection biology (Duneau et al., 2017), but presumably is a consequence of their different evolutionary histories.

Fungi cause the majority of insect diseases, infecting insects by directly penetrating the cuticle with a combination of cuticle-degrading enzymes and mechanical pressure (St. Leger, 2024; Wang & St. Leger, 2006). The entomopathogenic fungus *M. anisopliae* is widespread globally, and strain ARSEF549 (Ma549) is naturally pathogenic to *Drosophila (Lu et al., 2015)*. Unlike many other entomopathogens, it does not produce toxins that inhibit innate immune responses (Pal et al., 2007). Previous studies showed that about 9% of mutant *D. melanogaster* lines exhibit altered disease resistance to Ma549 (Lu et al., 2015) and that *Drosophila* Genetic Reference Panel (DGRP) flies from Raleigh, North Carolina, vary significantly in longevity after Ma549 infection (Wang et al., 2017). Our GWAS analysis of DGRP flies identified a complex network of immune and physiological genes, particularly those related to sleep, involved in *D. melanogaster* interactions with Ma549 (Wang et al., 2017). However, DGRP data only reflect genetic variation from a single location (North-Eastern USA), while a major portion of the genetic diversity of the species resides elsewhere, notably in Africa its ancestral home (Hutter et al., 2007). Thus, DGRP data are inadequate for studying global pathogen susceptibility.

Understanding the relationship between organisms and their habitats is central to ecological and environmental research. Diverse biomes, each with unique environmental conditions, exemplify this complexity (Kreft & Jetz, 2007). Examples of biomes with aseasonal climates include Tropical and Subtropical Moist Broadleaf Forests (TSMF), Tropical and Subtropical Grasslands, Savannas, and Shrublands (TSGSS) (Dinerstein et al., 2017), and Tropical and Subtropical Dry Broadleaf Forests (TSDF). In contrast, seasonal biomes like Temperate Broadleaf and Mixed Forests (TBMF), Mediterranean Forests, Woodlands, and Scrub (MFWS), and Desert Xeric Shrublands (DXS) experience greater climatic variability. Each biome’s unique climatic, topographic, and ecological variations significantly influence the resident organisms (Kreft & Jetz, 2007; Vasar et al., 2022), including the distribution of *D. melanogaster* populations and fungal species with pathogens being more diverse and numerous in the tropics (Tedersoo et al., 2014; Vasar et al., 2022; Větrovský et al., 2019).

We used Ma549 and 43 global *D. melanogaster* populations to investigate how environmental factors, genetic diversity, and sleep behavior affect disease resistance. The study focuses on temperature and humidity due to their significant impact on both fungal infection prevalence and insect resistance (Athanassiou et al., 2017). Increasing temperatures, changes in precipitation and extreme events under climate change threaten the persistence of many insects and their pathogens (St. Leger, 2021) so understanding adaptive responses to these stresses is crucial for predicting climate change impacts (Kellermann et al., 2018). We checked the relationship between abiotic and biotic stress using flies from the two ends of the resistance and biome spectrum. In particular, Monkey Hills (MH), St. Kitts, characterized by a humid and stable TSMF biome (Stancioff et al., 2018), and Ica, Peru (IP), a hyper-arid landscape with significant climatic variability (Salem, 1989).

Our findings show significant differences in Ma549 resistance among *Drosophila* populations from various global sites. This variation correlates with geographic origins and ecoregional conditions, suggesting persistent macro-physiological relationships between disease resistance and other traits. Sexual dimorphism also affects host-pathogen interactions regardless of geographic origin, with infection impacting sleep patterns more in males. Moreover, sexual activity influences disease resistance and sleep patterns, highlighting the complex interplay of genetic, environmental, and behavioral factors in shaping host responses to pathogens.

## Results

### The longevity of infected fruit flies varies by collection site and sex

We assessed the ability of 43 age-matched fly lines/populations from diverse geographic regions to survive infection by Ma549. Infection was performed via the natural route of applying spores to the body surface. Median lethal times (LT_50_) were monitored using five replicates (∼35 flies each) per sex per fly line, and the experiments were conducted at least twice. Lines with an unusually large disparity between males and females were screened three times to validate differences. A total of 14,957 male flies and 15,287 female flies were assayed, and the collection locations for the fly populations are listed in Table 1 and S1 Table.

**Table 1.**
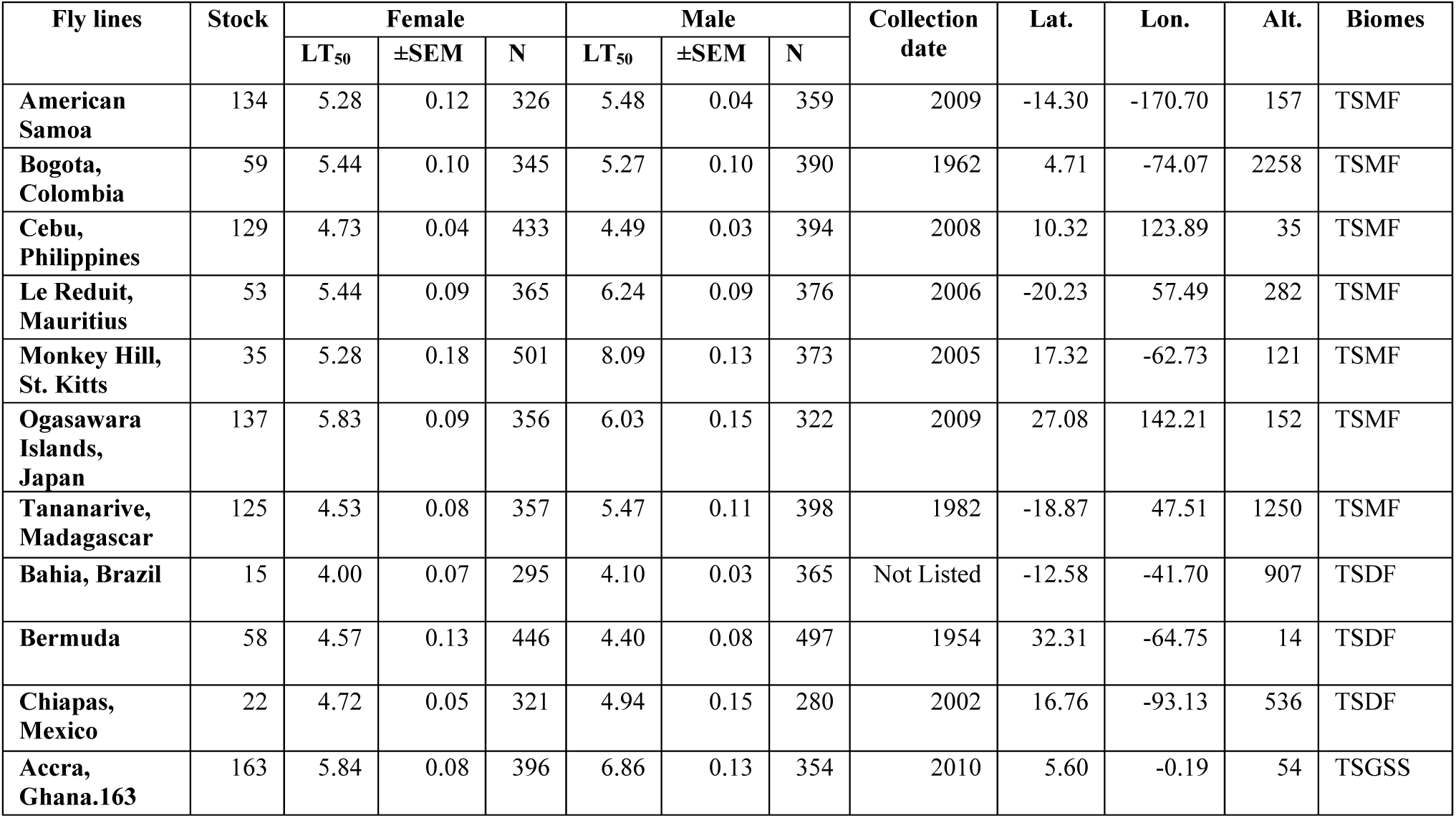
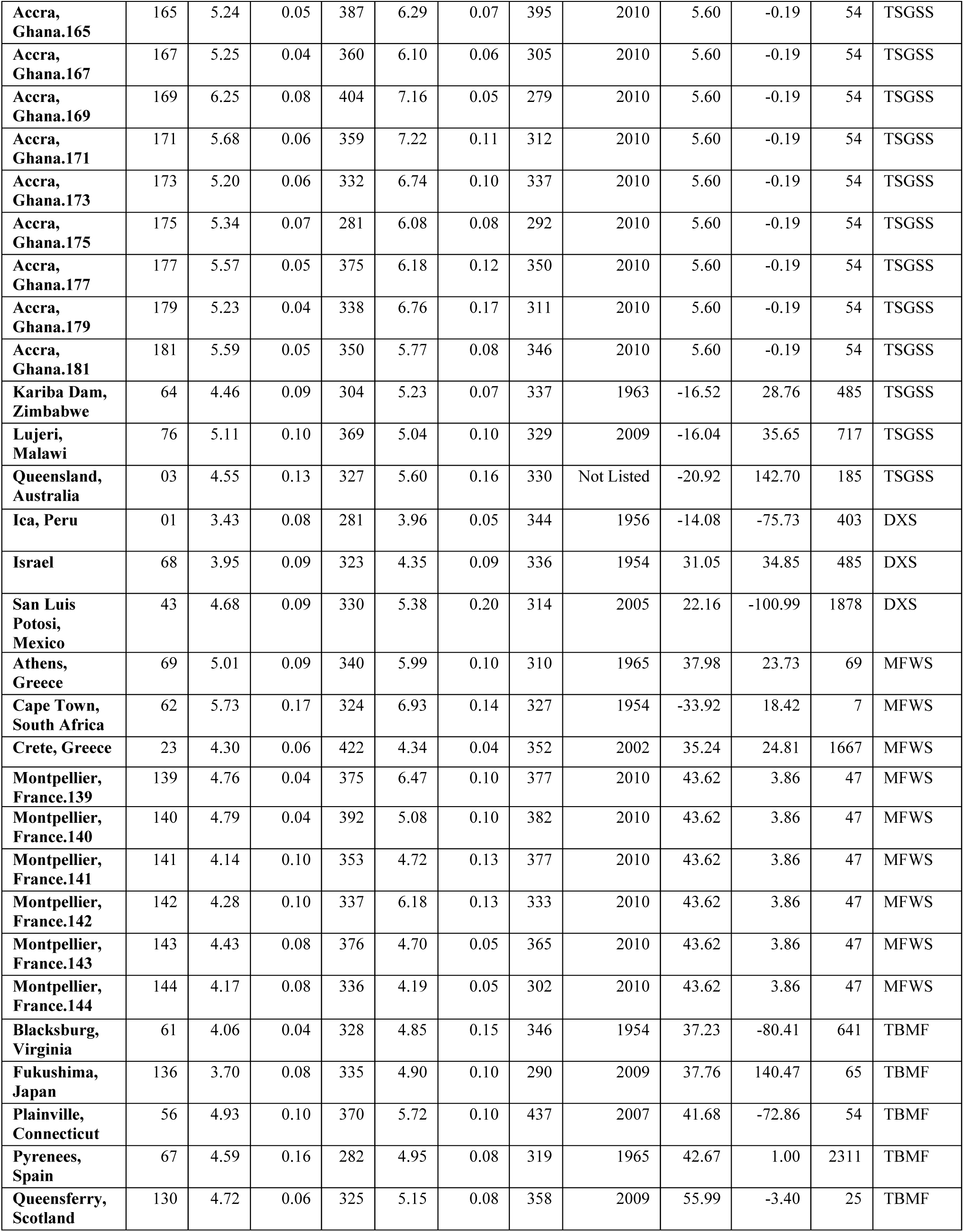
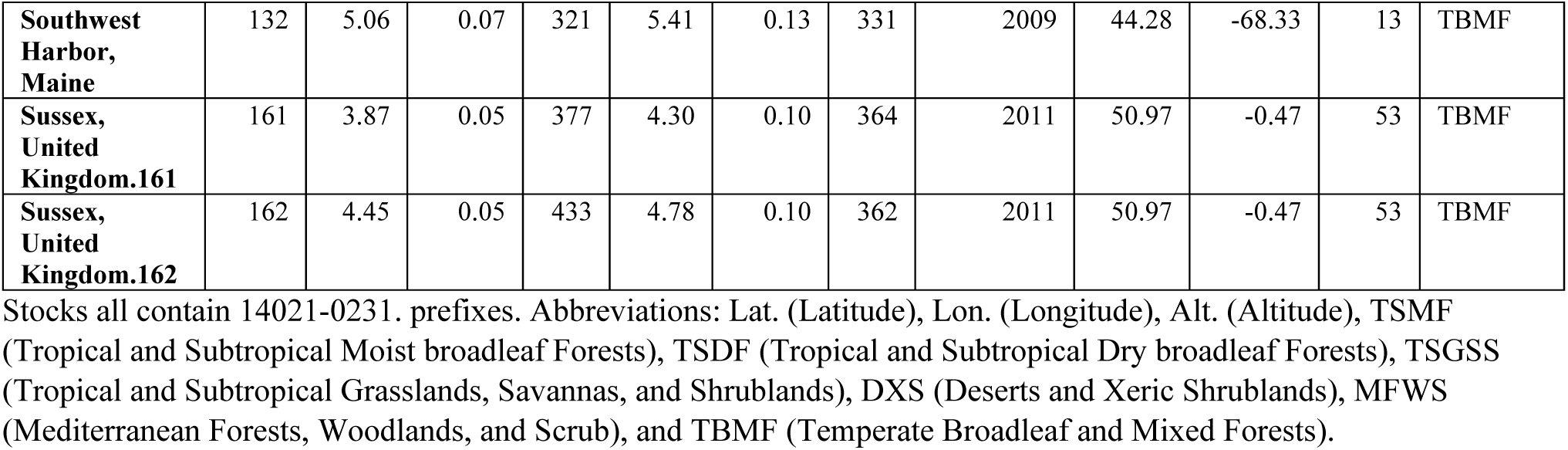
The localities, stock numbers, Ma549 LT_50_s, collection years, and geographic variables (including biomes) of 43 *Drosophila melanogaster* populations.

The 43 fly populations were sorted into six biomes based on the latitude and longitude of each collection site using ArcGIS Pro, and the 14 biomes defined in Ecoregions 2017 (Dinerstein et al., 2017) as follows: TSMF (Tropical and Subtropical Moist broadleaf Forests), TSDF (Tropical and Subtropical Dry broadleaf Forests), TSGSS (Tropical and Subtropical Grasslands, Savannas, and Shrublands), DXS (Deserts and Xeric Shrublands), MFWS (Mediterranean Forests, Woodlands, and Scrub), and TBMF (Temperate Broadleaf and Mixed Forests).

We also subdivided the 43 lines into four distinct geographical regions based on their collection sites: African (15), European/Middle East (13), Asian/Pacific (4), and American (11). The African lines were from Ghana (10), Madagascar (1), Malawi (1), Mauritius (1), South Africa (1), and Zimbabwe (1), spanning the TSMF, TSGSS, and MFWS biomes. The European/Middle East lines were from England (2), France (6), Greece (2), Israel (1), Scotland (1), and Spain (1), covering the DXS, MFWS, and TBMF biomes. The Asian/Pacific lines were from Australia (1), Japan (2), and the Philippines (1), representing the TSMF, TSGSS, and TBMF biomes. The American lines were from Bermuda (1), Brazil (1), Colombia (1), Mexico (2), Peru (1), and various locations in the United States (5), encompassing the TSMF, TSDF, DXS, and TBMF biomes (Table 1, Figure 1A).

**Figure 1.**
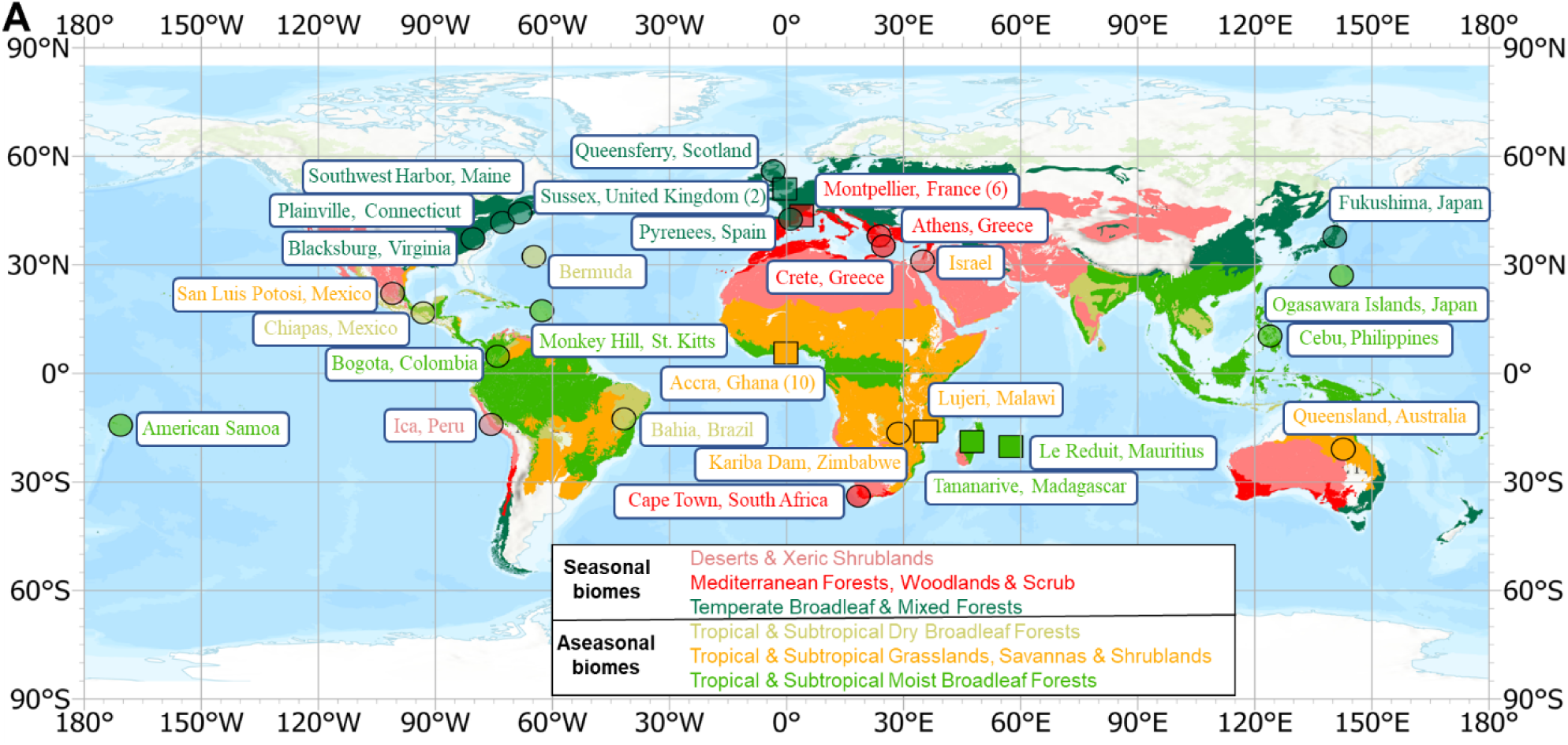
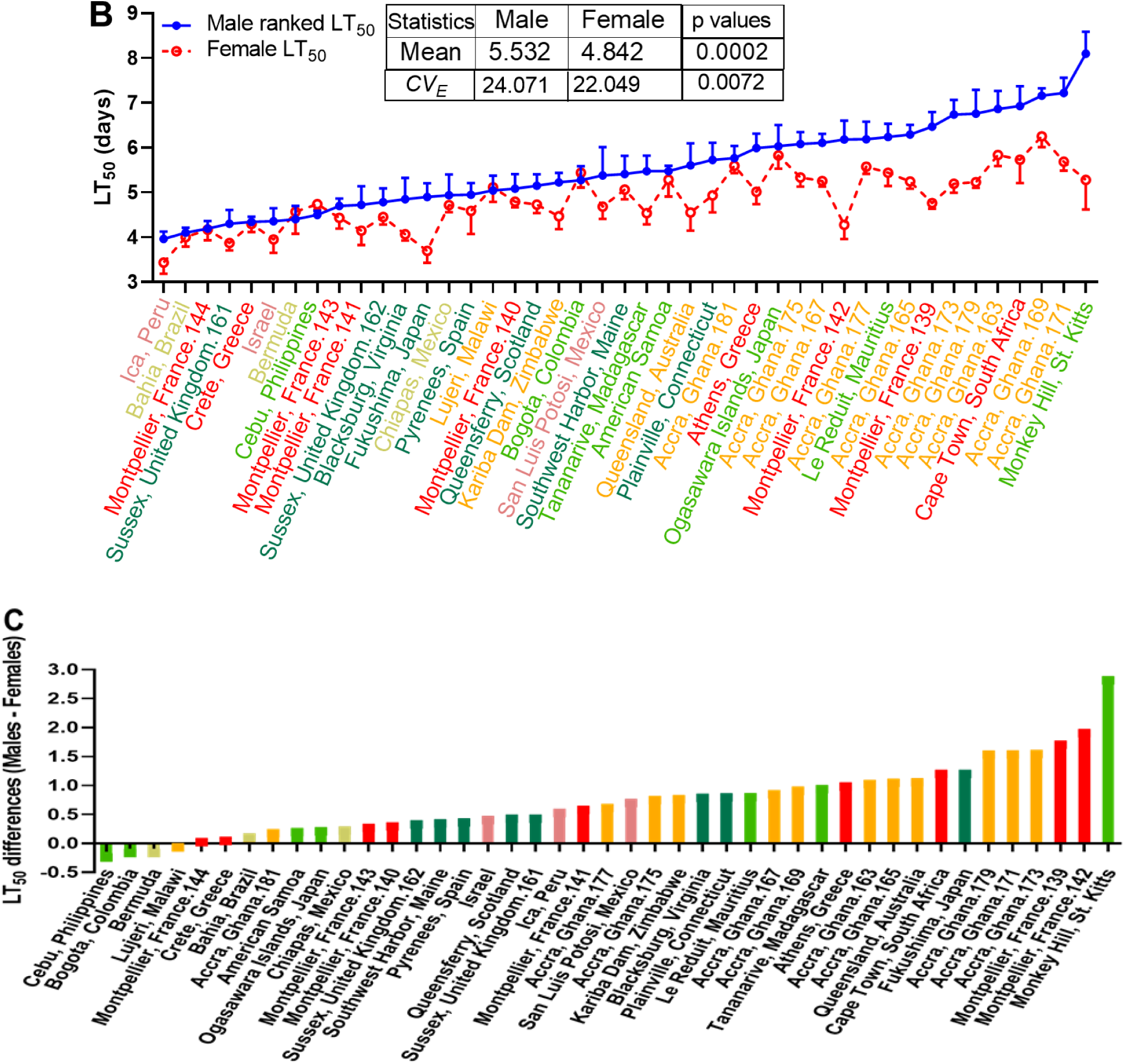
(A) Collection locations and biomes of *Drosophila melanogaster* populations. The geographic and biomes map (Dinerstein et al., 2017) generated by ArcGIS Pro displays the collection locations and biomes of 43 fly populations. The biomes are color coded for this and subsequent figures: Bright Green for Tropical & Subtropical Moist Broadleaf Forests (TSMF), Light Olive for Tropical & Subtropical Dry Broadleaf Forests (TSDF), Gold for Tropical & Subtropical Grasslands, Savannas & Shrublands (TSGSS), Pink for Deserts & Xeric Shrublands (DXS), Red for Mediterranean Forests, Woodlands & Scrub (MFWS), and Forest Green for Temperate Broadleaf & Mixed Forests (TBMF). The numbers in parentheses represent the number of fly lines collected at each site. The 22 circles mark the collection sites of previously deployed fly stocks(Brown et al., 2018) (gifted by Dr. Alex Keene), while six squares denote collection sites of 21 fly stocks from the *Drosophila* Species Stock Center. **(B)** The graph shows the mean LT_50_ values for females (red circles) and males (blue dots) across all 43 global fly lines, ranked by males from least to most resistant to Ma549 infection. Welch’s t test and unpaired t-test were used to obtain p-values for mean and coefficient of variation (*CV_E_*), respectively. **(C)** The graph illustrates sex-based disparities in Ma549 LT_50_ values, calculated by subtracting female LT_50_ values from male LT_50_ values for each of the 43 lines, averaged from 10-15 replicates (each containing an average of 35 flies), per sex per line in 2-3 experiments. Positive values indicate greater resistance in males, while negative values indicate greater resistance in females.

To identify sexual dimorphism, we measured resistance to Ma549 for males and females separately. Two-way ANOVA revealed significant interaction effects between line/genetic factor and sex (F_(42, 800)_ = 17.49, p < 0.0001), accounting for 9.87% of the total variation. Genetic factors were the most impactful (F_(42, 800)_ = 117.3, p < 0.0001), explaining 66.17% of the total variation, while sex contributed 12.81% (F_(1, 800)_ = 953.5, p < 0.0001) (Figure 2A, S1A and S8A Tables), showing that although males generally have longer lifespans, the extent of the differences between the sexes varies significantly across strains. Mean survival time (MST) and LT_50_ values showed strong correlations for both males (r = 0.9963, p < 0.0001) and females (r = 0.9992, p < 0.0001) (Figure 2B), indicating that both metrics reliably measure disease resistance. We mostly used LT_50_ values to compare resistance across lines and MST values for analyses related to plasticity.

**Figure 2.**
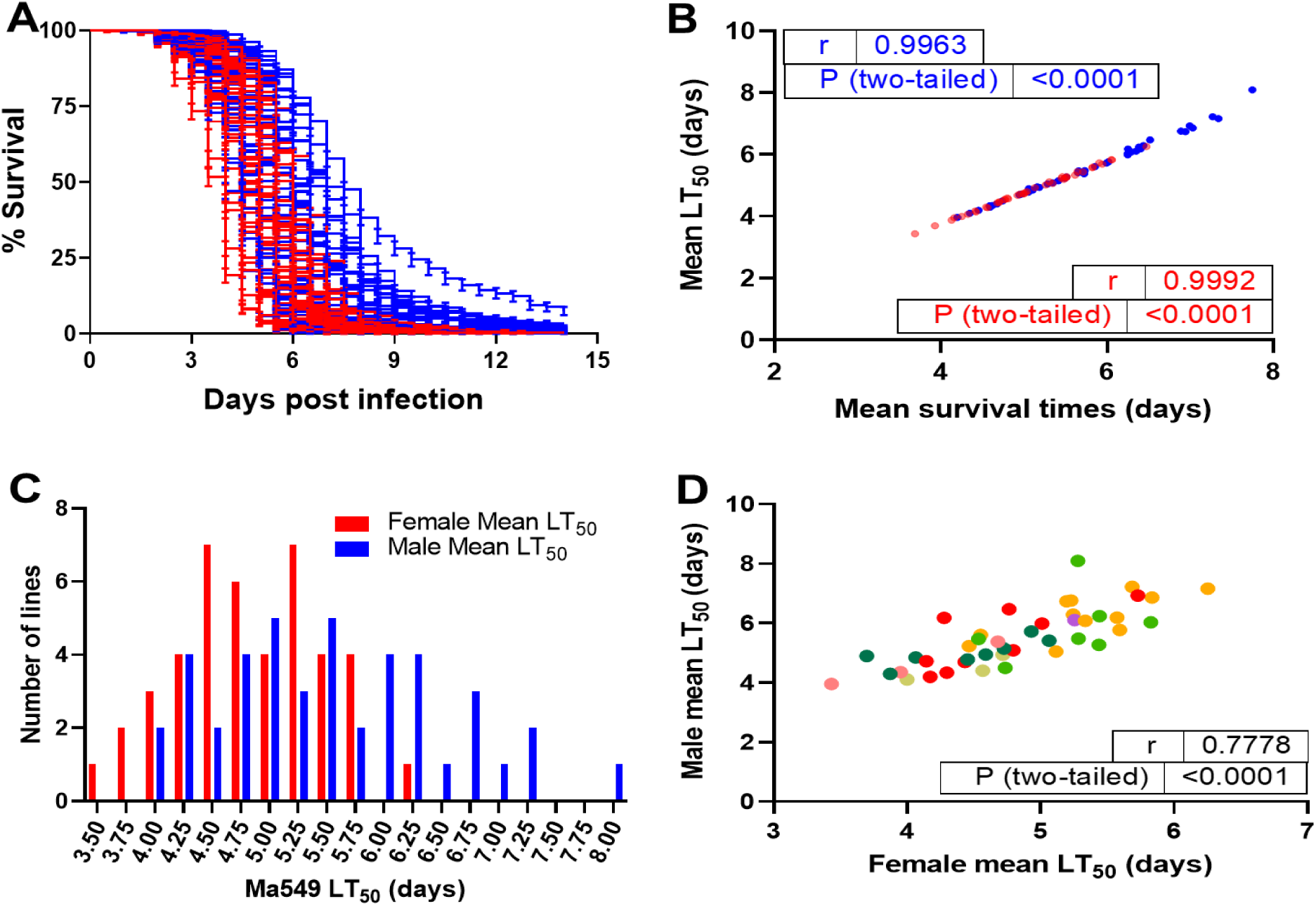
(A) The survival data of 43 *Drosophila* populations infected with Ma549. The mean percentage survival ± SEM for females (red, n = 281-501) and males (blue, n = 279-497) across 10-15 replicates per sex per line in 2-3 experiments. **(B)** The correlation between mean survival times and LT_50_ values in male (blue dots) and female (red dots) lines infected with Ma549, with Pearson correlation coefficient (r) and p-values indicated. **(C)** The distribution of longevity in females (red bars) and males (blue bars) confirming the normal distribution of mean LT_50_ values from the 43 lines using D’Agostino & Pearson and Shapiro-Wilk tests (p > 0.05). **(D)** Correlation analysis of LT_50_ values between the sexes, including the Pearson correlation coefficient (r) and p-values.

Kaplan-Meier survival plots confirmed a sex-based disparity in fly longevity, with males outliving females in 39 of 43 lines (S1A Table, Figure 2A and Figure 1B, C). The LT_50_ values for both female and male flies across diverse geographical regions were normally distributed (Figure 2C). The average LT_50_ for worldwide males (females) was 5.53 (4.84) days so males were, on average, 1.14-fold more resistant (t = 3.871, p = 0.0002, Figure 1B). The divergence between the most resistant and the most susceptible males (females) was 4.13 (2.82) days, ranging from 3.96 (3.43) to 8.09 (6.25) days (Figure 1B, C and Figure 2C, Table 1), showing that male flies displayed a wider range of LT_50_ values than females. However, cross-sex LT_50_ correlations were strong (r = 0.7778, p < 0.0001, Figure 2D), closely mirroring the DGRP lines (r = 0.7374, p < 0.0001) (Wang et al., 2017), indicating that many of the same genetic and environmental factors identified in the DGRP lines affect infection resistance in males and females worldwide.

The most susceptible line was from Ica, Peru (DXS biome), with an LT_50_ of 3.96 (3.43) days for males (females). The most resistant male flies were obtained from Monkey Hill (MH), St. Kitts (TMSF biome) with an LT_50_ of 8.09 days (Figure 1B, Table 1). Males from the MH populations exhibited 1.53-fold higher resistance than MH females, resulting in them having the largest sexual disparity of 2.81 days (Figure 1C, Table 1). Four lines (Cebu/Philippines, Bogota/Colombia, Bermuda, and Lujeri/Malawi) with intermediate LT_50_ values showed longer female lifespans (Figure 1C, Table 1), although these longevity differences were only significant in the Cebu/Philippines line (t = 3.68, p = 0.0003, Appendix 1—table 1). Notably, three of these four lines, excluding Lujeri/Malawi, originated from aseasonal regions characterized by high humidity and uniform temperatures. This suggests that females in these lines might have locally adapted to their specific environmental conditions differently than males did.

Ectothermic organisms commonly exhibit phenotypic plasticity, allowing a single genotype to produce different phenotypes to adapt to environmental stressors like temperature (Morgante et al., 2015). This plasticity is typically measured through variability among individuals (Morgante et al., 2015). Although the global lines are highly inbred they are not necessarily clones, so following common practice we estimated phenotypic plasticity by within-line variability expressed as a coefficient of environmental variation (*CV_E_*), which removes the association between mean and variance (David et al., 2005). The within-line standard deviation (*σ_E_*) has been used to measure variability among individuals in clonal lines of the DGRP (Morgante et al., 2015), and we used *σ_E_* to demonstrate that plasticity in disease resistance is highly heritable (Wang et al., 2017). *σ_E_* positively (p < 0.0001) correlates with *CV_E_* (males r = 0.7605; females r = 0.7157) and MST values (males r = 0.8300; females r = 0.5652), indicating that the most resistant lines also have the greatest variance (Figure 2—figure supplement 1A, B).

However, *CV_E_* is not significantly correlated with MST values in males (r = 0.2786, p = 0.0705) or females (r = - 0.1634, p = 0.2952) (Figure 2—figure supplement 1C), which is generally true for mean and *CV_E_* plasticity estimates in *D. melanogaster* (David et al., 2005). This suggests that disease resistance plasticity is independent of dimension, consistent with different genes controlling plasticity and MST values as previously found in the DGRP (Wang et al., 2017). The *CV_E_* ranged from 17.68 to 32.56 in males and 14.35 to 29.36 in females. Levene’s test was used to compare sex-based variations in disease resistance (Appendix 1—table 1). Of the 43 strains, 27 had higher *CV_E_* in males than females, with 23 showing significant differences per Levene’s test. Variation between sexes across the 43 lines was assessed by comparing *CV_E_* using a paired t-test, revealing significantly larger male variance across lines (t = 2.755, p = 0.0072, Appendix 1—table 1), confirming sex- and line-specific differences in variance. MH males showed the third highest plasticity and IP males the second lowest *CV_E_* (S1 Table 1, p ≤ 0.0024, Figure 2—figure supplement 1D, E), while for *σ_E_*, MH males were the most plastic and IP males the least (S1 Table 1, p < 0.0001, Figure 2—figure supplement 1F, G).

### Fly line and sex matter more than collection date

As the 43 fly lines had been inbred in laboratories for 13-70 years (Table 1), we examined whether domestication contributed to their variable longevity following infection. Excluding two lines with unspecified collection dates (Bahia/Brazil and Queensland/Australia), we analyzed the remaining 41 lines.

To assess the effect of fly line, sex, and collection date on fly longevity (LT_50_) following Ma549 infection, we used a linear mixed model formulated as: *Y_LT50_ = μ+S+C+S×C+L+ε*, where Y_LT50_ is the response variable representing LT_50_ values, μ is the overall mean, S denotes the fixed effect of sex, C represents the fixed effect of collection date, S×C indicates the interaction term between sex and collection date, L is the random effect of the fly lines, and *ε* is the residual error (Table 2).

**Table 2A.** Fixed effect tests on longevity effects of sex and collection date.

| Source | Nparm | DF | DFDen | F Ratio | p value |
| --- | --- | --- | --- | --- | --- |
| **Sex** | 1 | 1 | 41.79 | 20.9029 | <0.0001 |
| **Collection date** | 14 | 14 | 27.47 | 1.2574 | 0.2935 |
| **Sex\*Collection date** | 14 | 14 | 28.16 | 1.3107 | 0.2618 |

(Source: This is a list of the tested factors. Nparm: Number of parameters associated with the effect. DF: degrees of freedom for each source of variation. DFDen: Denominator degrees of freedom associated with the error term or residuals. F Ratio: the mean square of the factor divided by the mean square of the error. A p-value less than 0.05 means that the factor has a statistically significant effect.)

**Table 2B.** REML variance component estimates on longevity effects of fly lines.

| Random Effect | Variance Ratio | Variance Component | Std Error | 95% Lower | 95% Upper | p value | % of Total |
| --- | --- | --- | --- | --- | --- | --- | --- |
| **Fly lines** | 2.4757 | 0.4393 | 0.1457 | 0.1537 | 0.7245 | 0.0026 | 71.23 |
| **Residual** |  | 0.1775 | 0.0483 | 0.1109 | 0.3288 |  | 28.77 |
| **Total** |  | 0.6168 | 0.1457 | 0.4075 | 1.0423 |  | 100.00 |

(REML, restricted maximum likelihood, is a statistical method used to estimate variance components in linear mixed models, providing unbiased estimates by accounting for the loss of degrees of freedom when estimating fixed effects.)

The fixed effects model revealed significant main effects of sex (F_(1, 41.79)_ = 20.90, p < 0.0001) but not collection date (F_(14, 27.47)_ = 1.26, p = 0.2935) on LT_50_ values (Table 2A). There was also no significant interaction between sex and collection date (F_(14, 28.16)_ = 1.31, p = 0.2618). Thus, the significant effect of sex on LT_50_ values is independent of the collection date.

### Geography and biomes impact on host susceptibility to infection

Dividing the 43 global fly populations into four geographic regions (Africa, Europe/Middle East, Asia/Pacific, and America) and six biomes allowed us to determine the relative contributions of geographic location and biome characteristics in disease resistance. Stepwise regression analysis showed significant effects of both factors (p < 0.0001, Tables 1 and 3), with biome effects (η^2^ = 0.1291 or 12.91%) being about double those of geography (η^2^ = 0.0624 or 6.24%).

African populations, comprising 35% of the 43 lines, had mean LT_50_ values of 5.98 days (males) and 5.65 days (females), showing 1.12 to 1.24-fold greater disease resistance than other regions, including the N. Carolina DGRPs (Wang et al., 2017) (p ≤ 0.0484, Appendix 2—figure 1, and S8B Table). Males outlived females by 1.06 to 1.07-fold within each region (p < 0.0001), but African females outlived European/Middle Eastern males by 1.16-fold (p = 0.0021).

**Table 3.**
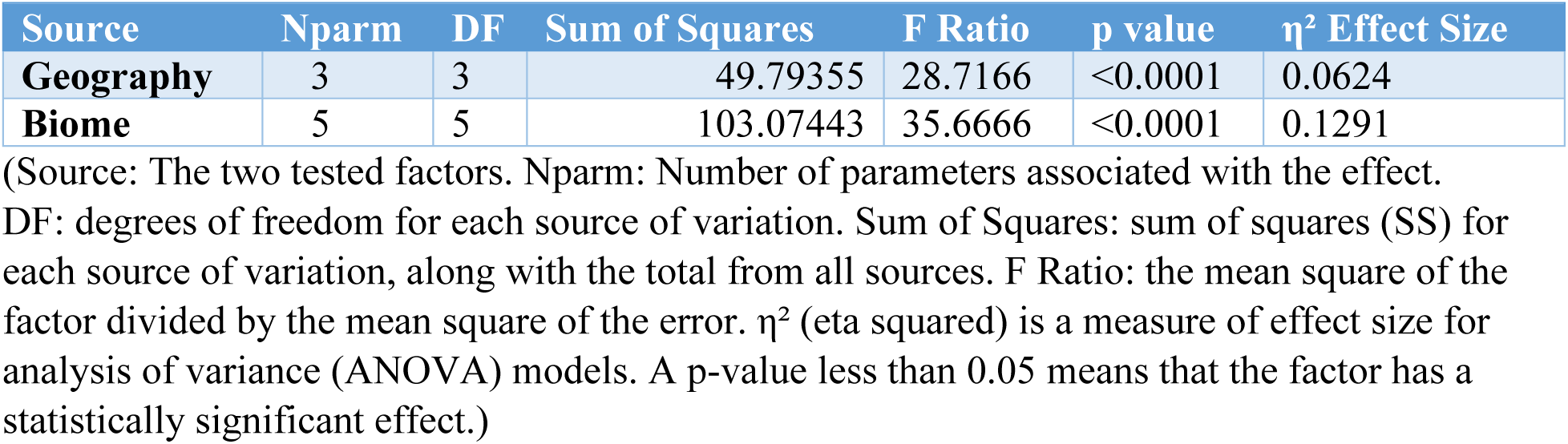
Impact of geography and biome on resistance levels.

| Source | Nparm | DF | Sum of Squares | F Ratio | p value | $\eta^2$ Effect Size |
| --- | --- | --- | --- | --- | --- | --- |
| **Geography** | 3 | 3 | 49.79355 | 28.7166 | <0.0001 | 0.0624 |
| **Biome** | 5 | 5 | 103.07443 | 35.6666 | <0.0001 | 0.1291 |

(Source: The two tested factors. Nparm: Number of parameters associated with the effect. DF: degrees of freedom for each source of variation. Sum of Squares: sum of squares (SS) for each source of variation, along with the total from all sources. F Ratio: the mean square of the factor divided by the mean square of the error. $\eta^2$ (eta squared) is a measure of effect size for analysis of variance (ANOVA) models. A p-value less than 0.05 means that the factor has a statistically significant effect.)

**Table 4.** Principal component loadings of environmental variables and LT50 for male and female flies.

| Principal component analysis (PCA) | Prin1 (57.5%) | Prin2 (15.8%) | Prin3 (10.9%) |
| --- | --- | --- | --- |
| T20-30°C & RH60-100% | 0.954346 | -0.030905 | 0.181506 |
| T20-30°C | 0.921278 | -0.241807 | 0.224949 |
| Average temperature (°C) | 0.837885 | -0.419570 | 0.265887 |
| Female $LT_{50}$ | 0.827146 | 0.214348 | -0.249415 |
| $\log_{10}$(latitude) | -0.826659 | 0.380707 | -0.131953 |
| Male $LT_{50}$ | 0.727170 | 0.141419 | -0.401798 |
| $\log_{10}$(precipitation) | 0.547000 | 0.669968 | 0.132934 |
| RH60-100% | 0.622824 | 0.636057 | -0.031578 |
| Dispersal distance | -0.354985 | 0.349947 | 0.751843 |

(Positive or negative loadings indicate a direct or inverse relationship with the principal component. Black or gray values denote strong or weak associations between LT<sub>50</sub> and certain environmental variables, respectively.)

### Environmental factors at the original collection sites influence host susceptibility to infection in the laboratory

We focused on temperature and humidity, key abiotic factors (Lotterhos et al., 2018) that act as stressors for many organisms, especially ectotherms when outside their preferred ranges (Bogaerts-Márquez et al., 2021). Latitude strongly correlates with annual temperature minimums and ranges, while altitude moderately correlates with annual relative humidity minimums and ranges (Appendix 3—figure 1A-F). Relative humidity decreases linearly with altitude (Peixoto & Oort, 1996), indicating altitude’s indirect influence on latitudinal humidity differences.

The clinal patterns in disease resistance for the 43 fly populations are shown in Appendix 3—figure 2A-G. Both females and males exhibit a marked and linear decrease in disease resistance (Ma549 LT_50_ values) with latitude (females: R² = 0.3342, p < 0.0001; males: R² = 0.2102, p = 0.0020). There was a smaller association between disease resistance and altitude in males (R² = 0.09931, p = 0.0396), that fell short of significance in females (R² = 0.07018, p = 0.0860). One potential explanation for a diversity of patterns in intraspecific analysis is that latitude or climatic variables *per se* may be less important in disease resistance than a population’s position relative to the species range (Woods et al., 2012). We found that disease resistance was greater towards the range center (Ghana) but the association was less than latitude, and populations from low latitudes, such as the Monkey Hill (MH) population were persistently more resistant despite distance.

Both sexes exhibit increased Ma549 LT_50_ values with annual average, high, and low temperatures at collection sites (S2 Table, Appendix 3—figure 2C-E). Annual mean precipitation (M) and precipitation in the wettest month (W) also correlate with LT_50_ values (S2 Table, Appendix 3—figure 2F, G, and Materials and Methods). Despite differences in intercepts caused by males being more resistant, regression slopes for males were slightly, but not significantly, steeper, suggesting the sexes were similar in how LT_50_ values trended with changes in temperature and precipitation.

The principal component analysis (PCA) summarized in Table 4 shows that the first principal component, capturing 57.5% of variance in disease resistance, reflects a general gradient of environmental temperature, latitude, and precipitation as well as relative humidity. The second component, explaining 15.8% of the variance, has relatively low correlations with male and female LT_50_ values. The third component, accounting for 10.9% of the variance, defines the impact of dispersal distance on male disease resistance. According to the eigenvalue greater than 1.0 Kaiser-Guttman rule, only the first three principal components (Prin 1-3) with eigenvalues greater than 1.0 are retained, as they explain more variance than any individual original variable.

### Abiotic and biotic stresses link ecological pressures to variations in disease resistance

While annual temperatures, precipitation levels, and wettest month precipitation provide a general environmental overview, they may not capture short-term variations or extremes that ectotherms must endure (Enriquez et al., 2020; Manenti et al., 2021). These variations will presumably be most pronounced in seasonal environments. Aseasonal biomes (TSMF, TSGSS, TSDF) typically have stable temperatures, high humidity, and abundant resources, whereas seasonal biomes (TBMF, MFWS, DXS) often experience greater environmental stresses. Aseasonal males (females) outlived seasonal males (females) by 1.15 (1.16)-fold (Figure 3A and S8C-D Table), and the most resistant lines from Monkey Hill (MH) and Africa were in the climactically uniform TSMF and TSGSS biomes.

**Figure 3.**
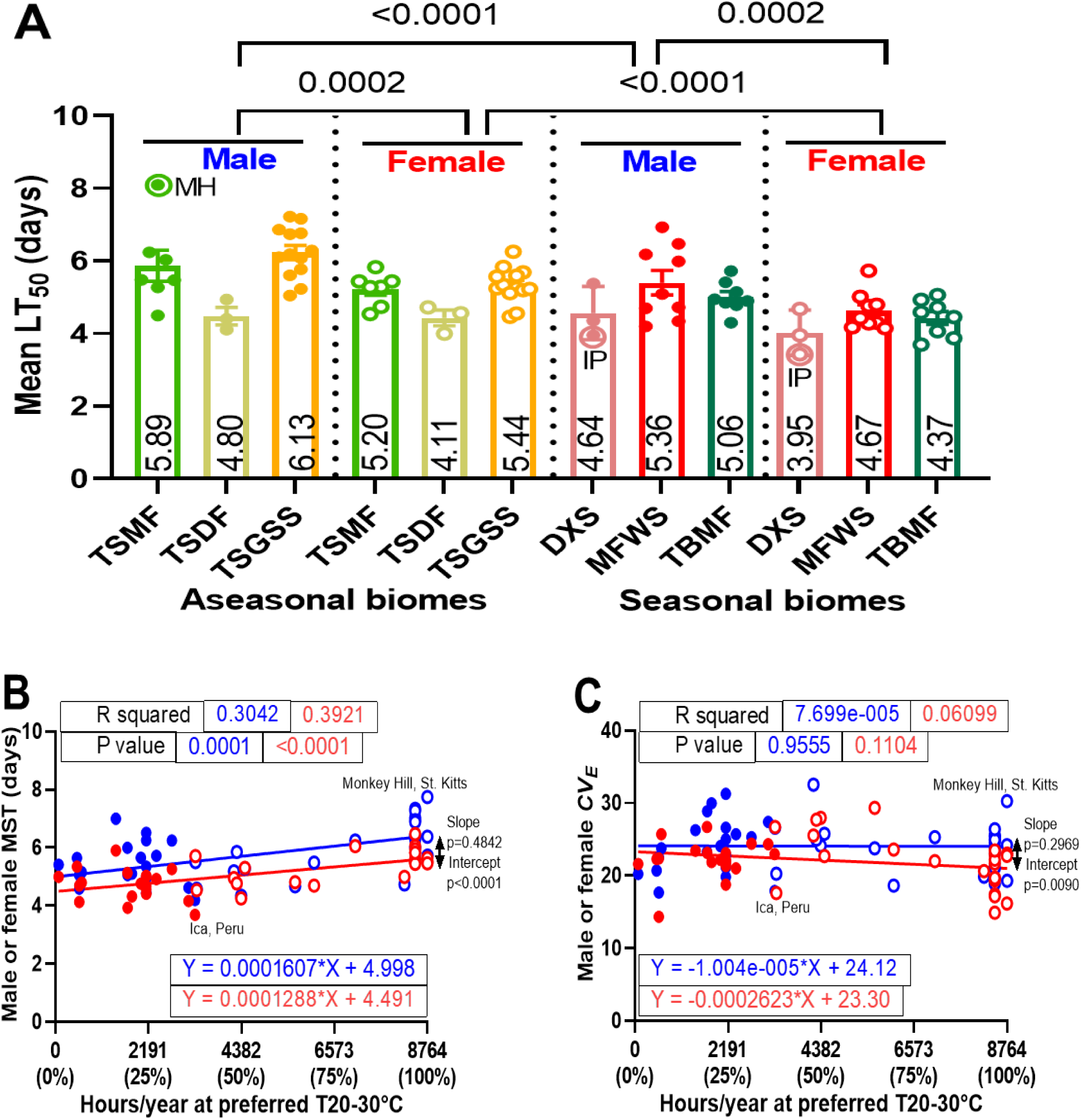
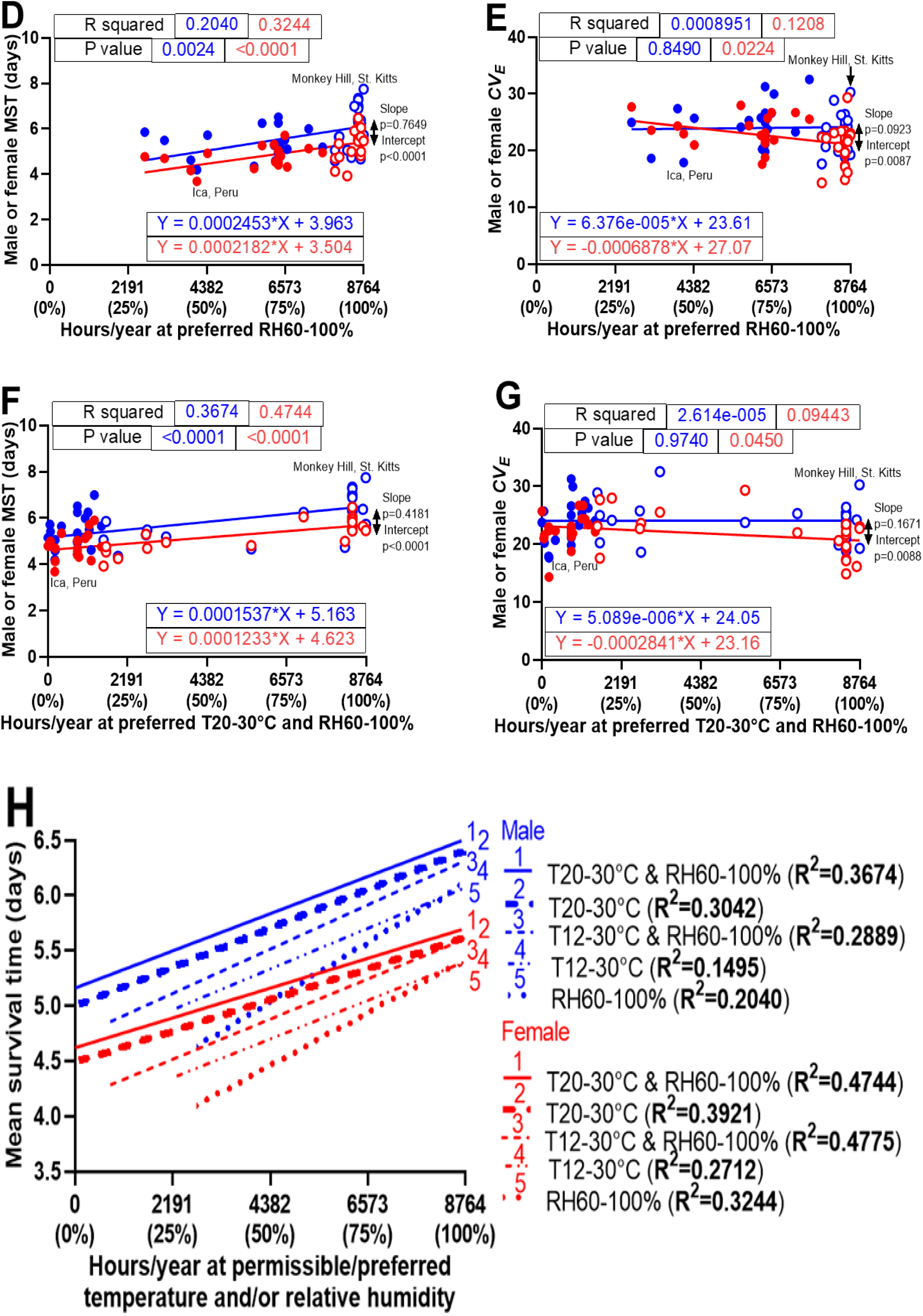
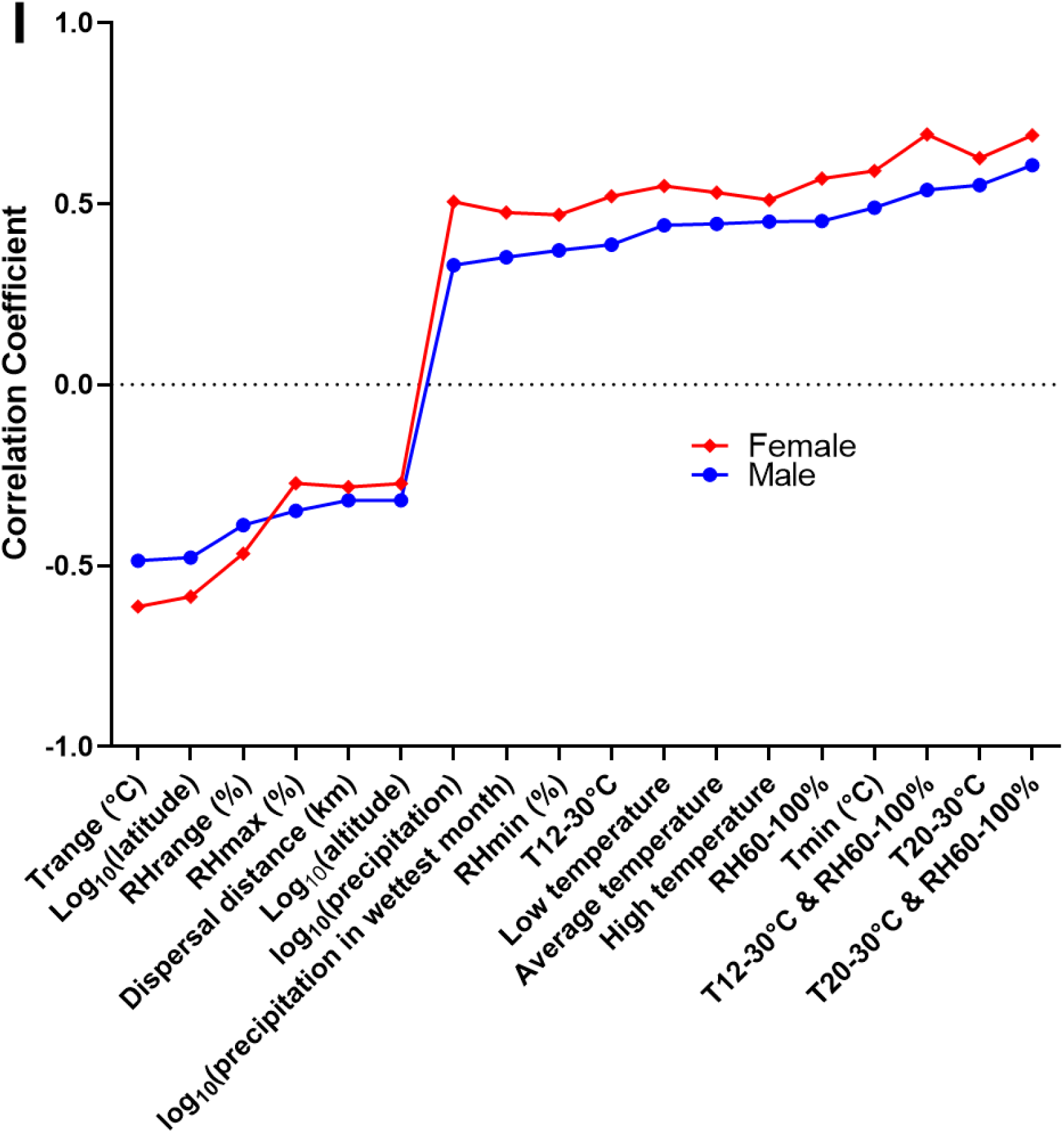
Biome variation in LT_50_ values among 43 global fly lines infected with Ma549 (**A**). The biome abbreviations and colors for the 43 worldwide fly lines match those in Figure 1A. LT_50_ values varied significantly across biomes and sexes, without a significant biome-sex interaction, as revealed by a two-way ANOVA. Significant p-values and LSM ± SEM from Tukey’s multiple comparison tests are shown. Males of the highly resistant MH line in the TSMF biome and highly susceptible IP lines in the DXS biome are indicated. Results for preferred 20-30°C temperature range with MST (**B**) and *CV_E_* (**C**), preferred humidity with MST (**D**) and *CV_E_* (**E**), and the combination of preferred temperature and humidity with MST (**F**) and *CV_E_* (**G**), including R^2^ values, p-values, and regression equations with slopes. Male (female) regression lines are blue (red). Aseasonal flies are depicted as circles, and seasonal flies as solid dots; this distinction is consistent in all figures. (**H**) Comparison of regression slopes (R^2^). (**I**) Correlation coefficients (r) between environmental factors and disease resistance (MST) in male and female *Drosophila* populations included, from left to right, annual temperature range (Trange), latitude, annual relative humidity range (RHrange), annual precipitation, precipitation in the wettest month, annual minimum relative humidity (RHmin), hours/year at 12-30°C (T12-30°C), low (night) temperature, average temperature, high (day) temperature, hours/year at 60-100% relative humidity (RH60-100%), annual minimum temperature (Tmin), hours/year of T12-30°C and RH60-100%, hours/year at 20-30°C (T20-30°C), and hours/year of T20-30°C and RH60-100%.

Understanding biome-specific factors may reveal how abiotic and biotic stresses link ecological pressures to variations in resistance Temperatures from 12-30°C and/or relative humidity above 60% are required for *D. melanogaster* development and fertility (Al-Saffar et al., 1996; Chakir et al., 2002). The optimal temperature for activity and development is 25°C (Hamada et al., 2008; Sayeed & Benzer, 1996), and fluctuating temperatures have adverse effects on lifespan (Siddiqui & Barlow, 1972), so we assumed a narrower 20-30°C temperature range would be preferred over the permissible 12-30°C. To investigate the influence of time spent outside these thresholds on disease resistance, the average annual exposure (hours per year) within the permissible temperature (12-30°C) and preferred humidity ranges (60-100%) was calculated using NASA Power Data. We analyzed 52,584 hourly temperature and relative humidity data points at the 28 collection sites from 2005 to 2010, corresponding to most of the 43 fly lines collection dates (S3 Table). The hours spent per year at 12-30°C ranged from 2,263 to 8,764 hours/year (Figure 3—figure supplement 1A-C). The hours spent per year at 60-100% humidity varied from 2,653-8,764 hours/year (Figure 3D, E). Linear regression analyses of the 43 fly populations revealed strong correlations for both sexes between disease resistance (MST values) and annual hours at 12-30°C or 60-100% humidity, showing a trend for increasing disease resistance as climatic uniformity increases. Hours within the preferred 20-30°C range varied from 75 to 8,764 hours/year across collection sites (Figure 3B, C and Figure 3—figure supplement 1G). The correlations with disease resistance are similar to those for the 12-30°C range but with increased R² values.

Annual hours between 12-30°C that coincided with 60-100% humidity spanned 742 to 8,764 hours/year across sites (Figure 3—figure supplement 1D-F), while annual hours at 20-30°C and 60-100% humidity fluctuated more, ranging from 1 to 8,764 hours (Figure 3F-G and Figure 3—figure supplement 1I). Linear regression analyses of the 43 fly populations revealed that disease resistance for both sexes was more strongly correlated with the annual hours at 20-30°C and 60-100% humidity than with 12-30°C and 60-100% humidity, or with temperature and humidity considered separately. No correlations were observed between *CV_E_* values and increasing climatic uniformity in males, while a negative relationship was found between *CV_E_* values and annual hours at 60-100% humidity in females with (Figure 3G) and without (Figure 3E) considering annual time at 20-30°C, suggesting that favorably uniform humidity conditions do not select for plasticity in females.

We next analyzed populations from the two ends of the resistance and biome spectrum to check the relationship between abiotic and biotic stresses. The males of the Monkey Hill, St. Kitts (MH) population show the highest disease resistance, whereas males and females of the Ica, Peru (IP) population are the most susceptible (Figures 2D and 4A, S1A-B Table). MH males (females) demonstrate a 2.04 (1.54)-fold greater resistance to Ma549 infection than IP males (females) (p < 0.0001, Figure 4A). Relative to the mean LT_50_ values across strains for males (females) of 5.53 (4.84), MH males (females) were approximately 1.46 (1.08)-fold more resistant and IP flies of both sexes 1.4-fold less resistant.

**Figure 4.**
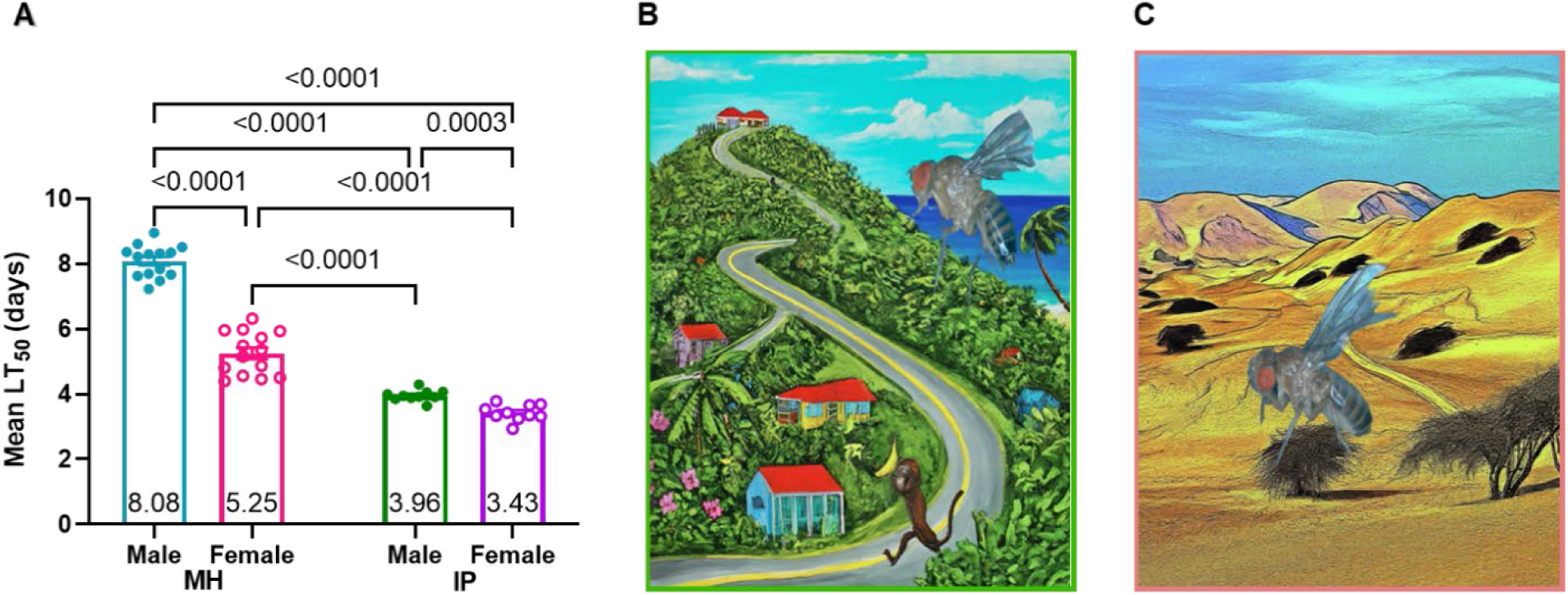
Sexual dimorphism and longevity of infected Monkey Hill, St. Kitts (MH) and Ica, Peru (IP) fruit flies **(A):** The data presents the mean ± SEM LT_50_ values across 14 replicates, with 501 (373) MH females (males) from 3 experiments, and 10 replicates with 281 (339) IP females (males) from 2 experiments. p-values are derived from Welch’s ANOVA tests with Dunnett’s multiple comparisons. Cartoon drawings of Monkey Hills, St. Kitts **(B)** and Ica, Peru **(C),** contrast their distinct biomes, illustrating the landscapes shaped by each location’s unique climatic environment.

MH in the TSMF biome (Figure 4B) has dense vegetation and a uniform maritime climate, whereas IP in the DXS biome (Figure 4C) experiences arid conditions and sparse desert-adapted vegetation (Table 1, and S2-3 Tables). Using Lomb-Scargle periodograms (Figure 4— figure supplement 1A-B, and S3B Table), we confirmed that from 2005 to 2010, MH maintained near-ideal conditions (20-30°C temperature, 60-100% humidity) for fly development 24 hours a day. In contrast, IP mostly experienced arid and fluctuating conditions with preferred temperature and humidity conditions coinciding only 2.12% of the time (Figure 4—figure supplement 1C-J).

We subjected male and female MH and IP flies to a desiccating environment to determine whether there might be trade-offs with disease resistance. Kaplan-Meier analyses demonstrated that IP males (females) were 2.00 (1.53)-fold more resistant to desiccation stress than MH males (females) (p < 0.0001, Figure 5A-B). Furthermore, females of both lines were more desiccation resistant than males, with IP and MH females outliving their male counterparts by 1.14 and 1.49-fold, respectively (p ≤ 0.0033, Figure 5C-D) (S4A Table). These differences between IP and MH appear adaptive as only IP evolved in a dry environment; however, they totally reverse the direction of resistance to Ma549.

**Figure 5.**
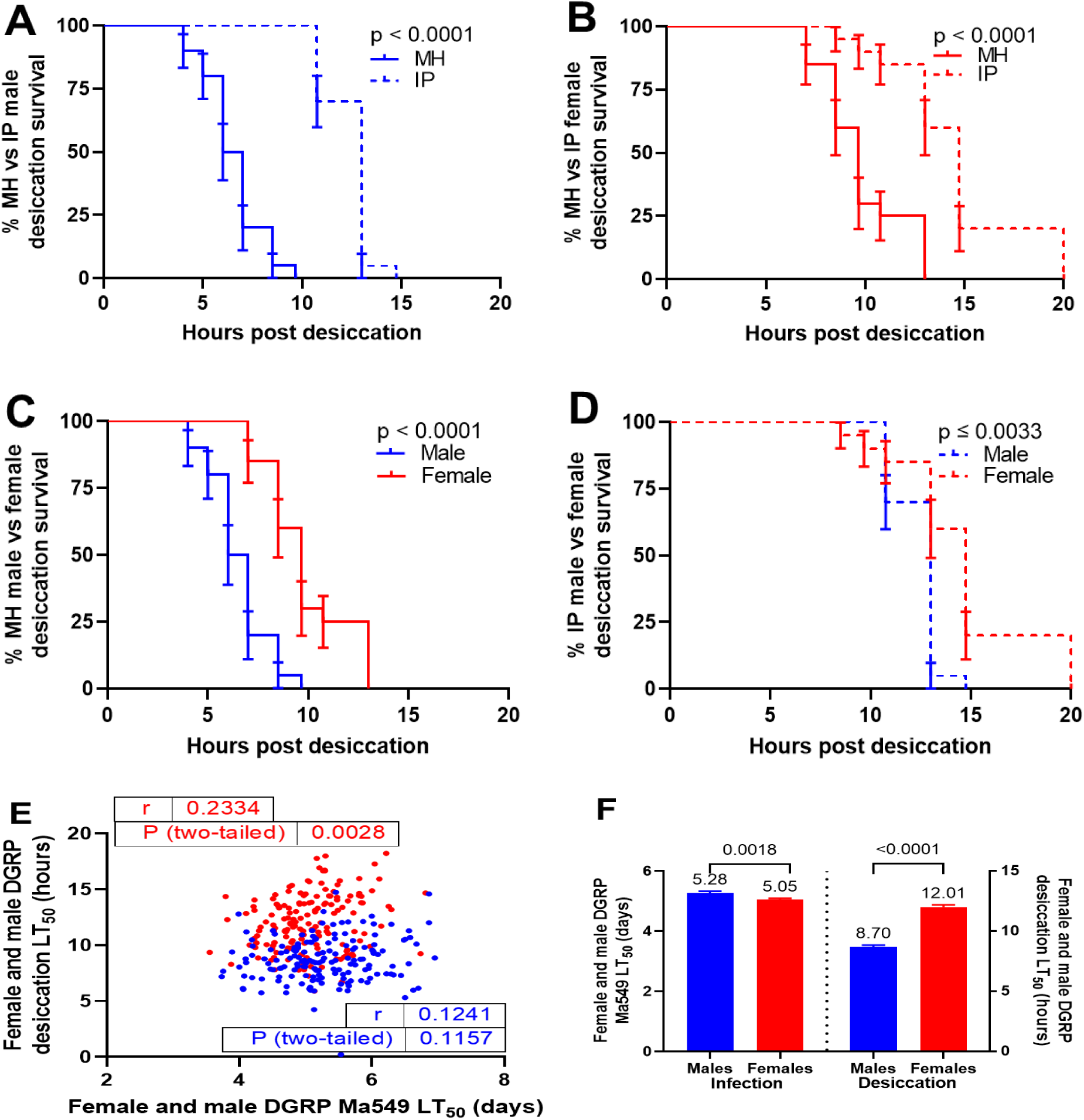
Differences in desiccation resistance. Kaplan-Meier plots illustrate survival differences between MH and IP lines **(A, B)**, and between sexes **(C, D).** MH males (females) from five replicates (10 flies each) across two trials are shown in turquoise (crimson), and IP males (females) in green (purple). Statistical significance was assessed using log-rank and Wilcoxon tests. Correlation analysis between desiccation and Ma549 LT_50_ values of 162 DGRP females and males **(E)**, with Pearson correlation coefficients (r) and p-values indicated. Males are represented by blue dots and females by red dots. Sexual dimorphism in resistance to disease (left y-axis) and desiccation stress (right y-axis) for the 162 DGRP fly lines **(F)** is represented as mean ± SEM LT_50_ values from disease resistance (Wang et al., 2017) and desiccation resistance data (Rajpurohit et al., 2018). p-values are derived from Welch’s t-tests.

To examine the relationship between disease resistance and desiccation in a larger sample, we reanalyzed disease resistance in the DGRPs (Wang et al., 2017), and the desiccation resistance trait quantified in a later study of the DGRPs (Rajpurohit et al., 2018) (S4B Table). We found that resistance of DGRPs to desiccation significantly correlates with female resistance to Ma549 (r = 0.2334, p = 0.0028), but not males (r = 0.1241, p = 0.1157) (Figure 5E). Moreover, similar to MH and IP, these DGRP lines exhibited a complete reversal in the direction of sexual dimorphism in these two resistance traits: males outlived females by 1.05-fold in Ma549 resistance, whereas females outlived males by 1.38-fold in desiccation resistance (Figure 5F).

### Correlation between sleep and disease resistance

Using healthy females from 22 global populations, Brown et al., (2018) (Brown et al., 2018) demonstrated that higher monthly average temperatures at collection sites correlate with increased sleep duration in the laboratory. Our current study found a correlation between collection site temperatures and longevity post-Ma549 infection, while our previous DGRP analysis linked disease resistance with several sleep parameters (Wang et al., 2017) suggesting an avenue to explore the relationship between sleep indices and disease resistance in a global population.

Using the sleep behavior data provided for females from 22 global populations (Brown et al., 2018), we identified significant positive correlations between LT_50_ values and both 24h sleep duration (r = 0.5586, p = 0.0069, Figure 6A) and sleep bout length (r = 0.6230, p = 0.0020, Figure 6B), alongside a negative correlation with sleep bout number (r = -0.4703, p = 0.0272, Figure 6C). These results underscore a significant link between healthy consolidated sleep patterns and Ma549 infection outcomes.

**Figure 6.**
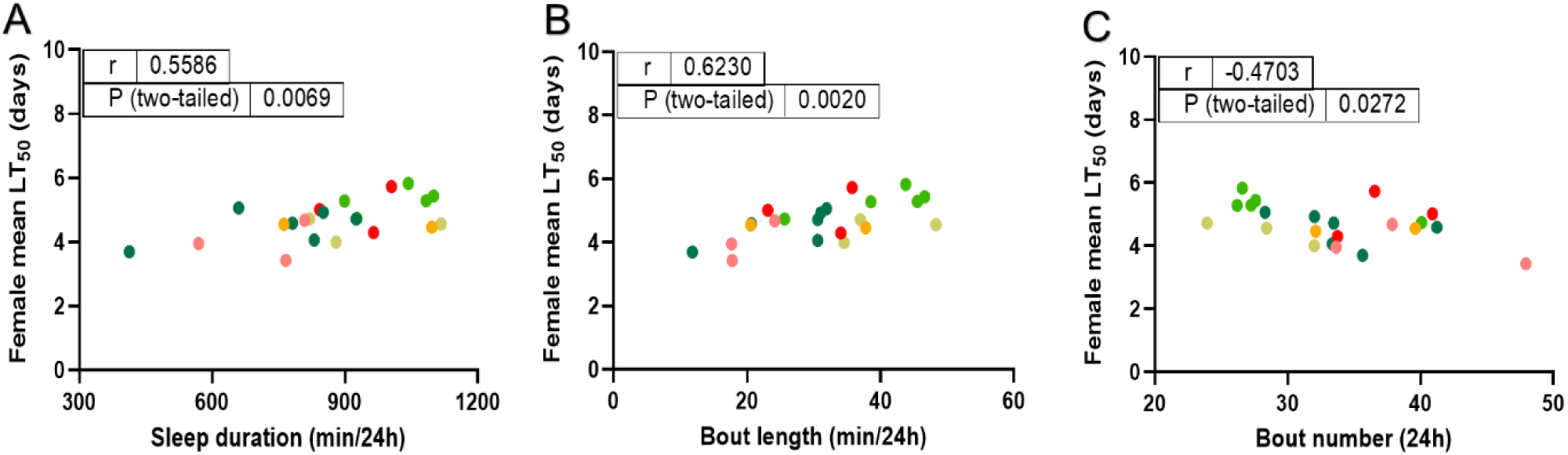
Correlations of LT_50_ values among Ma549-infected females and publicly available sleep indices for 22 of the lines (Brown et al., 2018) deployed in our study. LT_50_ values are positively correlated with sleep duration (**A**) and bout length (**B**), and negatively correlated with bout number (**C**). Individual Pearson correlation coefficients (r) and p-values are indicated in the figures.

### Characterization of sickness sleep triggered by Ma549 infection

The significant correlation between healthy female daily sleep patterns and Ma549 infection outcomes could be because latitudinal and climatic gradients have acted independently on sleep and disease resistance or alternatively be because sleep and resistance are evolutionarily intertwined. We investigated post-infection sleep behaviors in MH (resistant) and IP (susceptible) flies using the *Drosophila* Activity Monitor (DAM2) (Shaw et al., 2000) focusing on the first two days post-infection before the onset of overt symptoms. We also investigated a MFWS population from Crete, Greece (CG). Male (female) CG flies have LT_50_ values of 4.34 (4.30) days (Table 1), with only 0.04-day difference between the sexes (Figures 2D and 9A).

The sleep data (S5 Table) showed that uninfected mated MH females had 16% less nighttime sleep, but 36% more daytime sleep than uninfected males (p < 0.0001, Figure 7A-D), which averaged out to females sleeping about 12 minutes less in a day. Ma549 infection did not disrupt the circadian rhythm in MH flies, but significantly increased daytime sickness sleep (extra sleep following infection as defined by Davis & Raizen (2017) (Davis & Raizen, 2017)) compared to uninfected controls (p < 0.0001, Figure 7E-H). Mated males showed the largest increase in daytime sleep (107%), followed by virgin males (68%) (p < 0.0001, Figure 7F, H). Infected mated (virgin) females showed only a 17% (14%) increase compared to uninfected females (p < 0.0001, Figure 7E, G). Nighttime sleep in females was unaffected by infection (p ≥ 0.0661, Figure 7E, G) but increased by 2% in mated and virgin males (p ≤ 0.0119, Figure 7F, H).

**Figure 7.**
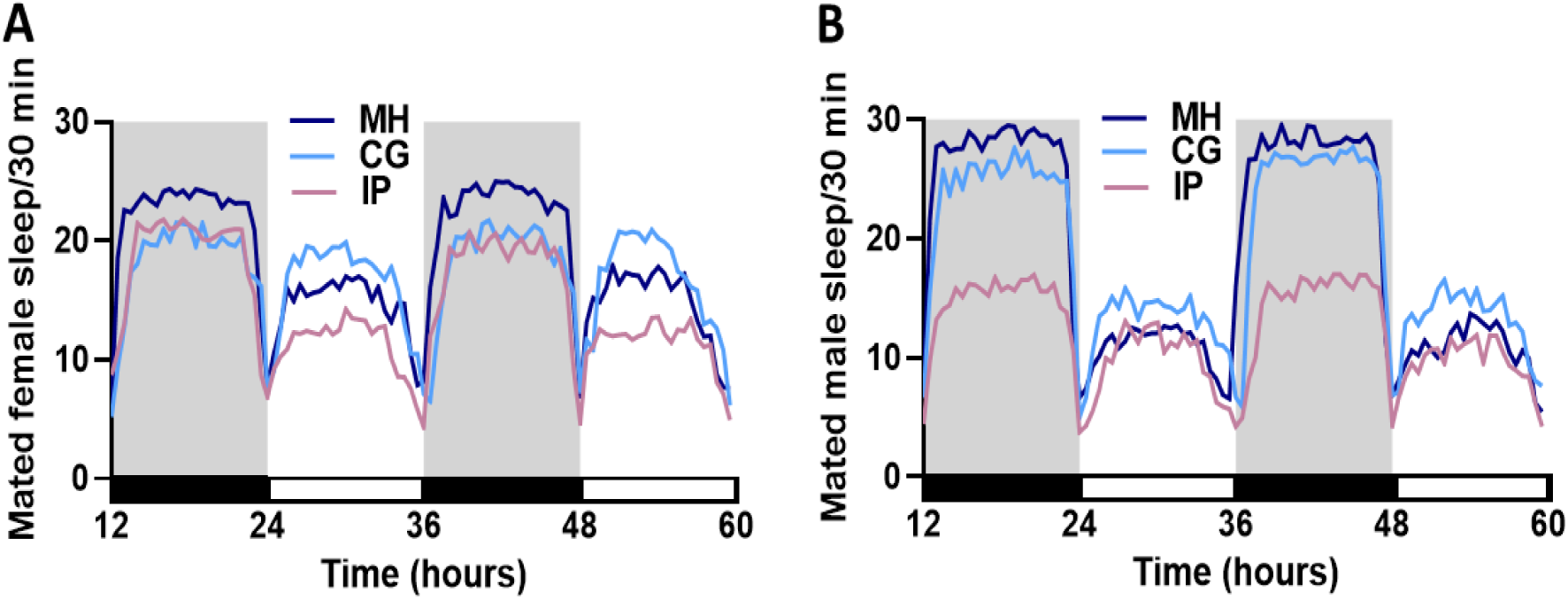
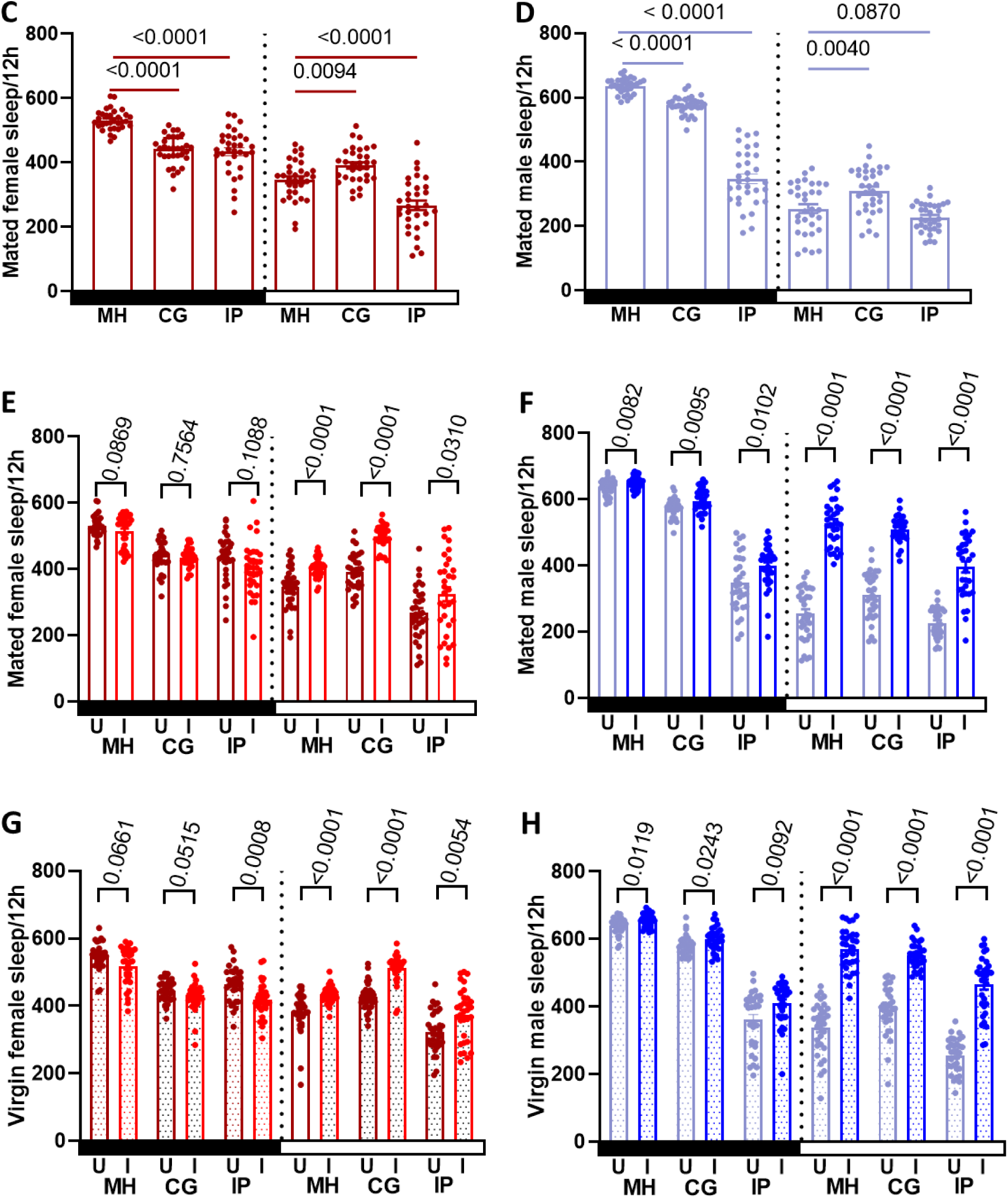
Ma549 infection affects sleep and survival. Sleep profiles depict sleep in min per 30 min from 12-60 h spanning two light-dark cycles for 32 flies pooled from three trials for uninfected mated MH, CG, and IP females **(A)** and males **(B)**, with gray shading indicating nighttime. Sleep comparisons were made between uninfected mated MH versus CG or IP females **(C)** or males **(D)**, and uninfected (U) versus infected (I) mated females **(E)**, mated males **(F)**, virgin females **(G)**, and virgin males **(H)** within the MH, CG and IP lines. Bars illustrate nighttime (black) and daytime (white) sleep for uninfected (dark red for females, sky blue for males) and infected (scarlet red for females, navy blue for males) mated and virgin (shaded) flies, with each data point representing the average sleep/12h of a single fly (n = 32) from 12-60 h post infection. In panels (**C)** and **(D)**, means ± SEM and p-values were obtained from one-way ANOVA. In panels (**E)-(H)**, Mann-Whitney U tests, Welch’s t-tests, and unpaired Student’s t-tests were used for various comparisons, with mean ± SEM and p-values indicated.

Compared to uninfected MH flies, CG females and males slept less at night and more during the day (Figure 7A-D). CG females slept 24% less at night and 26% more during the day than the males (p < 0.0001, Figure 7A-D). Infection with Ma549 increased daytime sleep: mated males by 64%, virgin males by 41%, mated females by 27%, and virgin females by 20% (p < 0.0001, Figure 7E-H). Similar to MH flies, nighttime sleep was minimally affected by Ma549 infection (Figure 7E-H).

The highly susceptible IP flies had shorter sleep durations than MH or CG flies with males being more active (slept less) at night (Figure 7A-D). Unlike MH and CG lines where males slept longer, IP females slept 25% longer at night (p < 0.0001, Figure 7A-D), and 18% longer during the day (p = 0.0172, Figure 7A-D). Ma549 infection increased daytime sleep in IP flies by 74% in mated males (p < 0.0001, Figure 7F), 81% in virgin males (p < 0.0001, Figure 7H), 21% in mated females (p = 0.0310, Figure 7E), and 16% in virgin females (p = 0.0054, Figure 7G), with minimal effects on nighttime sleep (Figure 7E-H).

As males are generally more resistant than females, we investigated sexual dimorphism in sleep as a factor in disease resistance (Figure 8A, B, S5 Table). Compared to females, Ma549 infection significantly increased daytime sleep in mated and virgin males of MH (30% and 31%), IP (22% and 24%), and virgin CG (8%) lines (p ≤ 0.0096). The effect was minimal in mated CG males (2%, p = 0.1862), as was the difference in disease resistance between mated CG males and females. Mated and virgin MH and CG males slept 27% and 40% longer at night, respectively (p < 0.0001), compared to females, while IP males did not differ significantly from females (p ≥ 0.7619). These sleep pattern differences aligned with disease resistance, with mated (virgin) males outliving females by 53.8% (51.2%), 0.95% (8.3%), and 15.4% (13.80%) in the MH, CG, and IP lines, respectively (p < 0.0001, Figure 8E, Table 1 and S1C Table), suggesting that, except for mated CG males, mated and virgin males exhibit longer sickness sleep as well as greater disease resistance than females following Ma549 infection.

**Figure 8.**
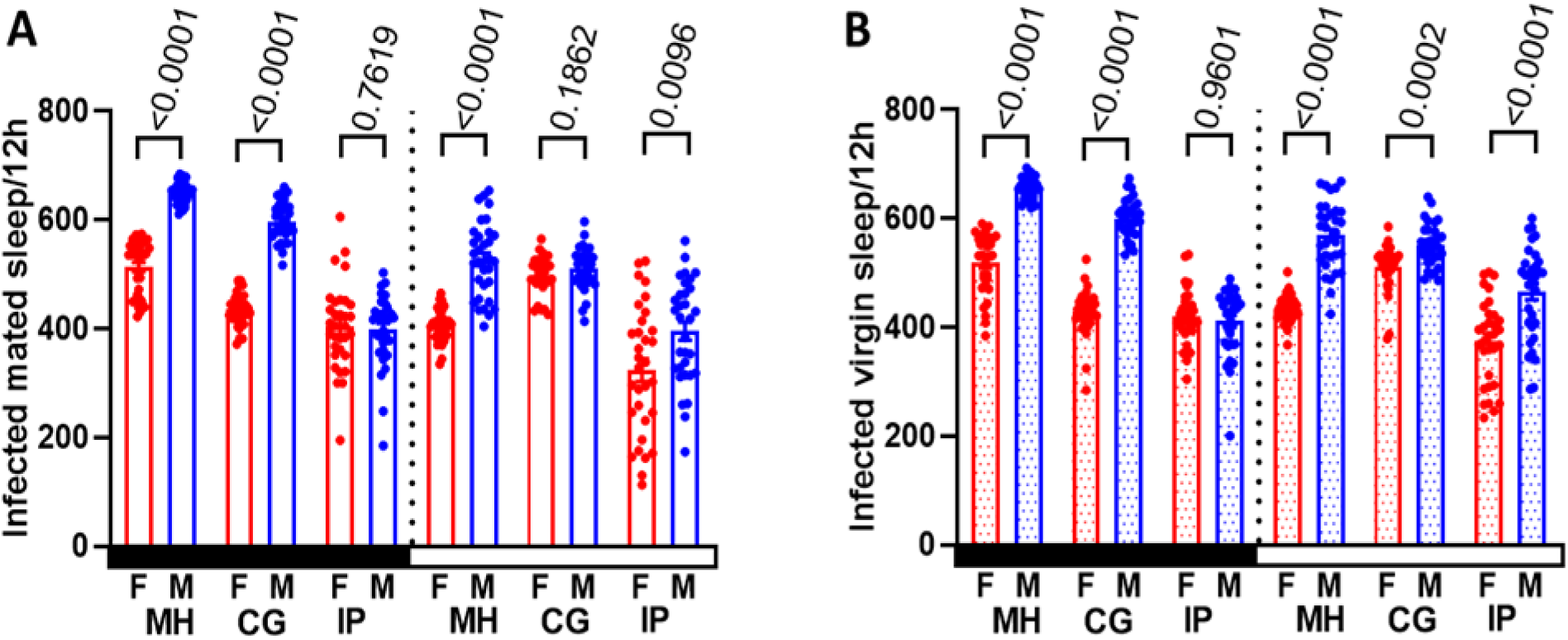
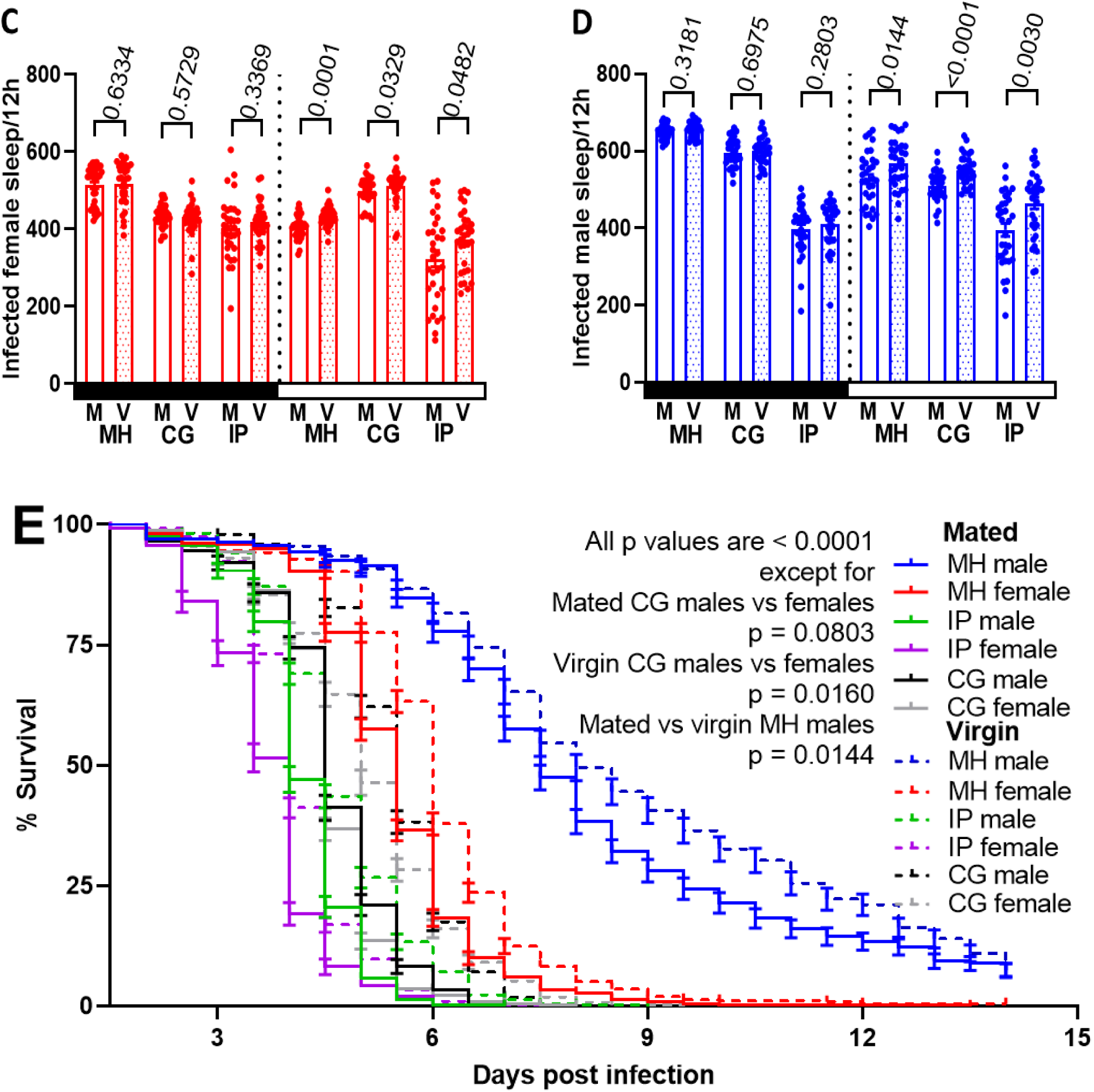
Sexual dimorphism in sleep was compared in mated **(A)** and virgin **(B)** MH, CG, and IP flies. Sleep was averaged per 12 h for each fly (n = 32) during day and night from 12-60 h post infection, pooled from three trials. Mann-Whitney U tests, Welch’s t-tests, and unpaired Student’s t-tests were used for various comparisons, with mean ± SEM and p-values indicated. Variations in nighttime and daytime sleep between mated and virgin infected female (**C**) and male (**D**) flies are shown, with each point representing a single fly’s average sleep per 12 h (n = 32 pooled from three trials) from 12-60 h post-infection. Significance was assessed between mated and virgin females or males within each lines using various tests, with mean ± SEM and p-values indicated. Black (white) rectangles on the x-axis represent nighttime (daytime), and shaded bars denote virgin flies. Line abbreviations: V for virgin and M for mated. Kaplan-Meier plots **(E)** depict survival differences between female (n = 281-585) and male (n = 337-434) or between mated (n = 281-501) and virgin (n = 337-585) flies in each line post-infection. Log-rank and Wilcoxon tests were used for statistical comparisons, with p-values indicated for general comparisons and specific group comparisons.

Mating status did not affect nighttime sleep of infected females or males across the three strains (p ≥ 0.2803, Figure 8C, D and S5C Table), but sexual activity significantly decreased daytime sickness sleep irrespective of sex and infection status (p ≤ 0.0482), with infection reducing daytime sleep by 3-15% in mated flies compared to virgins. Following infection, virgin females (males) exhibited 10.3-14.7% (8.5-20.2%) higher survival compared to mated counterparts (p < 0.0001, Figure 8E, Table 1 and S1C Table), highlighting the significant influence of sexual activity on sleep regulation and immunity in flies, regardless of genetic background or specific fly lines.

ANOVA was used to validate the effects of genetic line, sex, mating status, and infection on both nighttime and daytime sleep. The results are detailed in Appendix 4—table 1 (derived from data in S5B Table) and visualized in Figure 8—figure supplement 1A, B).

### The impact of host variation on fungal fitness

We analyzed the relationship between longevity and Ma549 colonization (measured as fungal load in colony forming units CFU) of the most resistant (MH), susceptible (IP), and representative average (CG) *D. melanogaster* lines. MH males (females) survived Ma549 infection 2.86 (2.28)-fold longer than CG males (females) (p ≤ 0.0006, Figure 9A), with no significant differences between CG males and females (p = 0.9897). Host variation significantly impacted Ma549 fitness, with CFUs appearing 2.0 days post-infection in IP flies, and 4.0- and 7.0-days post-infection in virgin MH females and males, respectively. Mated IP females were the least able to restrain fungal growth with 12,791 ± 700.05 CFUs/fly on day 3.5, while virgin MH males had the lowest counts at 16 ± 1.21 CFUs/fly on day 7.0 (Figure 9B-D, S6 Table). Fungal loads increased rapidly near fly death in all lines, indicating a breakdown of host resistance. Males, regardless of mating status, had lower fungal loads than females (Figure 9B-D), and virgin females delayed fungal proliferation by approximately 0.5-days compared to mated flies (Figure 9C-D), aligning with differences in their LT_50_ values (Figures 4A and 9A, Table 1 and S1C Table). Correlation analyses showed that LT_50_ values (Table 1 and S1C Table) were strongly correlated with both the onset (initial time point at which fungal growth becomes detectable) and peak times (time point maximum fungal growth achieved) of CFUs (r = 0.9910, p < 0.0001, Figure 9E-F) across mated and virgin IP, CG, and MH females and males.

**Figure 9.**
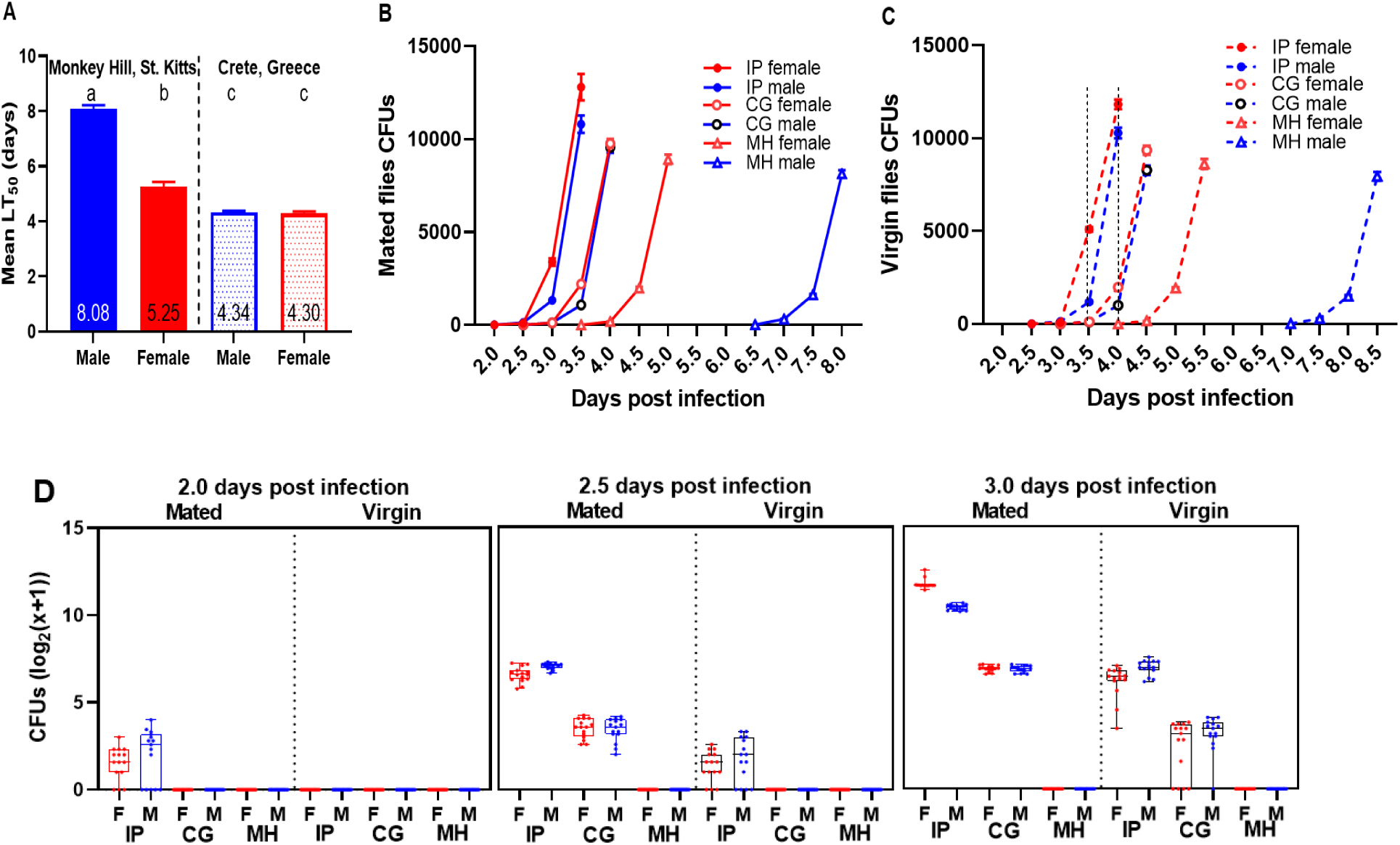
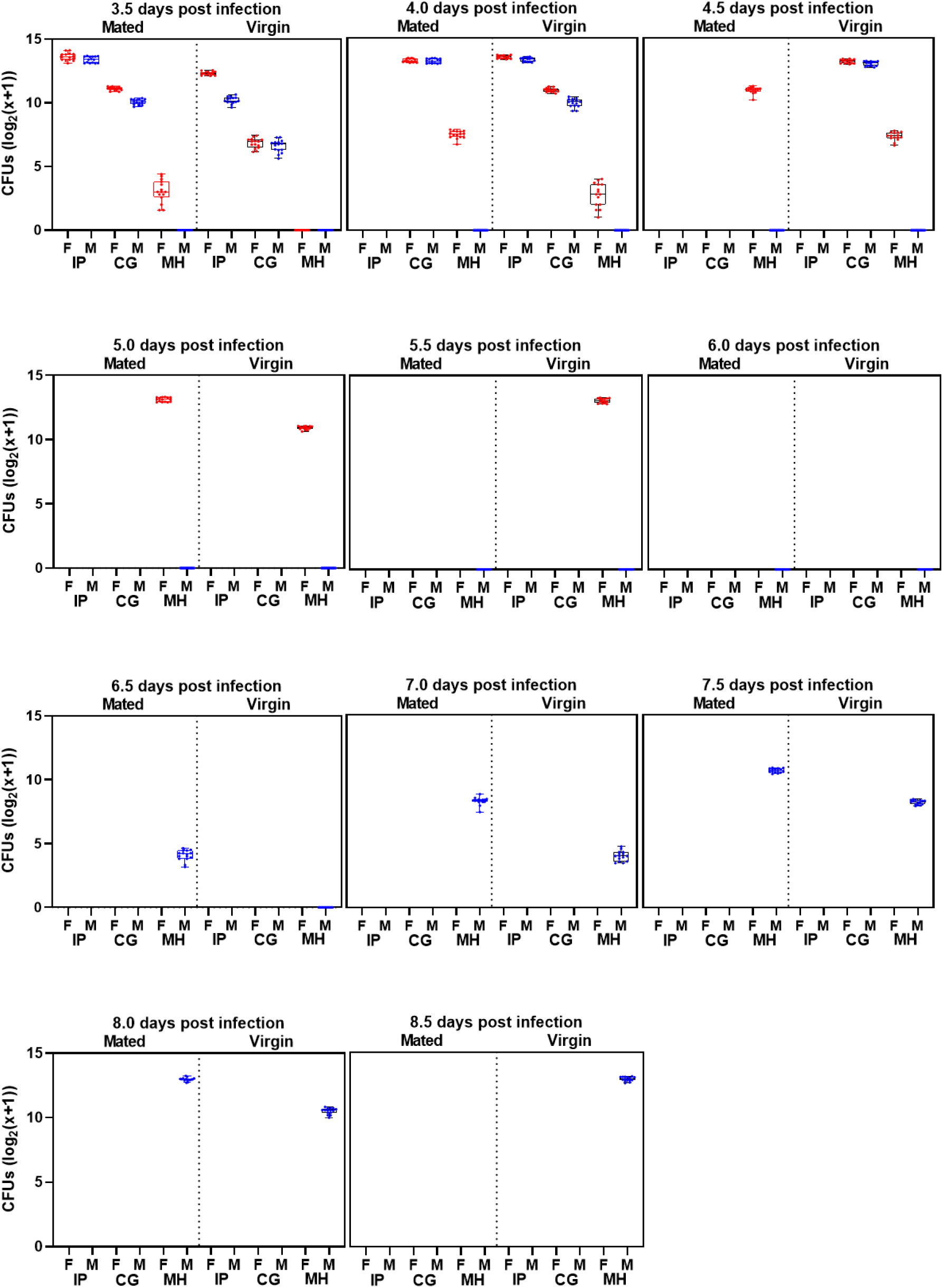
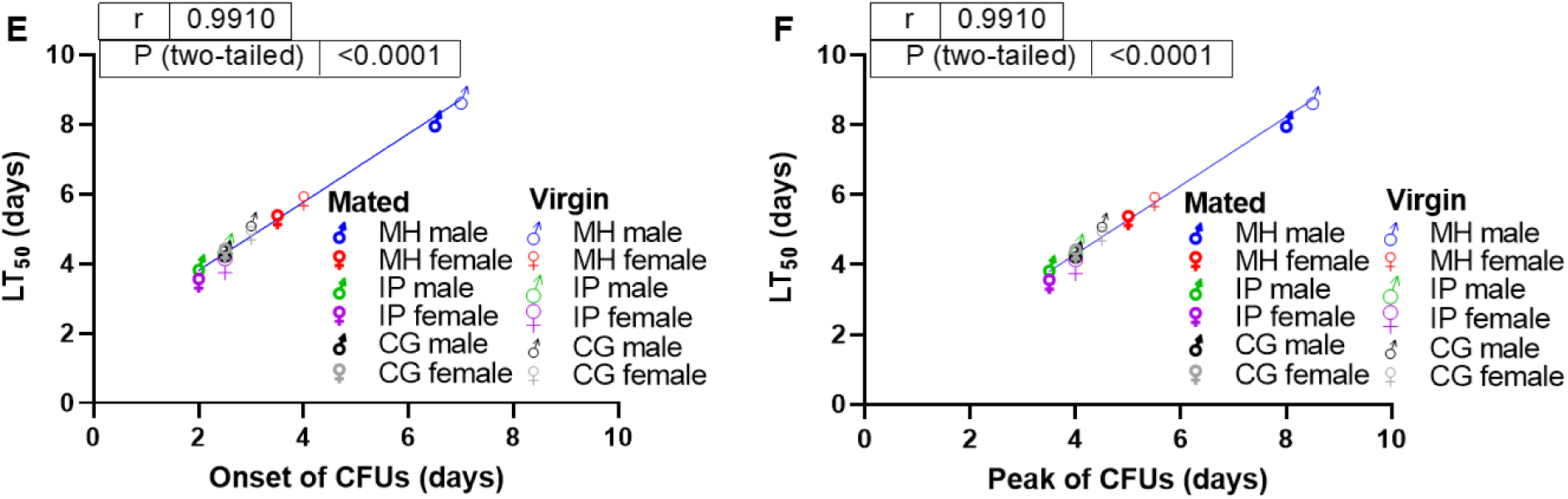
(A) Sexual dimorphism and longevity of infected Monkey Hill, St. Kitts (MH) and Crete, Greece (CG) fruit flies. The data display the mean ± SEM LT_50_ values from 14 replicates, with 501 MH females and 373 MH males across 3 experiments, and 422 CG females and 352 CG males across 2 experiments. Bars not connected by the same letter are significantly different. CFU production time course in mated **(B)** and virgin **(C)** CG, IP, and MH flies following Ma549 infection. Each data point represents the average of CFU values from 15 flies. Means ± SEM and p-values were obtained from two-way ANOVA. Time course of log-transformed Ma549 fungal loads in the hemolymph of CG, IP, and MH flies following Ma549 infection **(D).** Box plots depict each time point for each of the 15 individual replicates per sex, mating status, line, and time point across three repeated experiments, showing the mean ± SEM of CFUs. Correlations between LT_50_ values and CFU onset **(E)** or peak **(F)** were analyzed for infected mated and virgin CG, IP, and MH females (red) and males (blue). Pearson correlation coefficients (r) and p-values indicated.

These findings confirm that a fly’s line, sex and mating status all influence fungal invasiveness, with proliferation coinciding with and likely causing death due to an inability to tolerate high fungal levels (Figure 9).

### Potential adaptive differences between populations from Africa, France and Raleigh, North Carolina

Based on the observed longevity differences across geographic regions and biomes, we explored the underlying genetic basis by analyzing single nucleotide polymorphisms (SNPs) linked to disease resistance. Using SNPs from the North American DGRP (Wang et al., 2017), we investigated allele frequency differences that could indicate local adaptation to pathogens in the DGRP, African and French isolates (S7 Table). We previously found that most polymorphisms associated with Ma549 resistant DGRP lines were in the lower range of the allele frequency spectrum, with frequencies below 0.2 for 41% of the genes. These low-frequency alleles had larger effects on LT_50_ values than common alleles, resulting in the most Ma549 resistant DGRP lines having a preponderance of these alleles (Wang et al., 2017). Given that African lines typically exhibit higher resistance than DGRP lines and are localized in an aseasonal biome (Appendix 2—figure 1), we hypothesized that these alleles may be more prevalent in Africa.

We defined subsets of rare SNPs (S7 Table) previously linked with quantitative traits (LT_50_ values or CV_E_, micro-environmental plasticity as described (Morgante et al., 2015)) in Ma549 resistant DGRP lines using their corresponding positions in African and European populations. We used the PopFly database (Hervas et al., 2017) to download genome sequences from African lines corresponding to the 10-bp region flanking each SNP. Besides the 10 lines from Ghana, we included the Siavonga, Zambia (ZI) population, which consists of 207 individuals (large sample size) and is the most diverse among all sampled populations but with minimal admixture from non-African populations (Hervas et al., 2017; Lack et al., 2016).

The African populations exhibited higher frequencies of 11 out of 24 alleles affecting LT_50_ values in the DGRP lines. Variants typically showed modest 2-to 3-fold differences between African and American populations, but some SNPs displayed approximately 10-fold differences, mostly favoring African populations, resulting in a 22.3% higher average frequency of resistance alleles in African than American lines (Figure 10A). Sixteen out of 29 alleles influencing CV_E_ values were more prevalent in African lines, with frequency differences leading to twice as many CV_E_ variants for high plasticity in African genomes than in the DGRP lines (Figure 10B). Low frequency (<0.08) DGRP variants in *Rab26* (exocrine secretion), *Xpd* (DNA repair, UV damage) and at an intergenic site were the major alleles (>0.5) in African populations, whereas DGRP variants in two poorly understood genes, *CG32066* (DGRP frequency 0.1311) and *CG13229* (DGRP frequency 0.2327), were absent in Zambia and Ghana populations. An extended search in the PopFly database across additional African populations (136 Cameroon, 680 Ethiopia, 136 Gabon, 176 Kenya, 88 Malawi, 88 Nigeria, 136 Rwanda, 504 South Africa, 176 Uganda and 88 Zimbabwe) revealed the DGRP SNP variant of *CG32066* in single flies from South Africa (SD39N) and Uganda (UG19), but the DGRP *CG13229* variant was not found in Africa.

**Figure 10.**
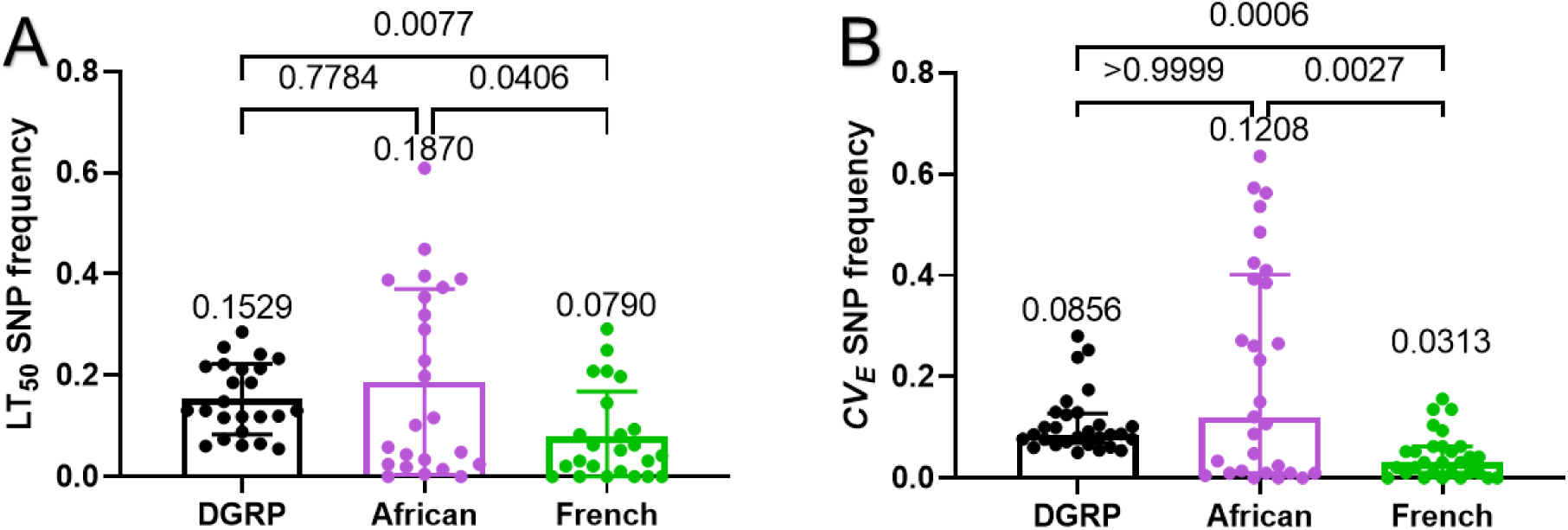
Frequency analysis of single-nucleotide polymorphism (SNP) for LT_50_ **(A)** and coefficient of variation (*CV_E_*) **(B)** among DGRP, African, and French populations. Welch’s ANOVA tests (W_(2, 42.24)_ = 6.289, p = 0.0041) reveled significant differences in LT_50_ SNP frequency across the DGRP, African, and French groups. Dunnett’s multiple comparisons test identified significant p-values and mean ± SEM (A). The Kruskal-Wallis test showed significant differences (H = 16.58, p = 0.0003) among the groups, with significant p-values and median ± IQR from Dunn’s multiple comparisons test noted (B).

The DGRP is an admixture of African and European populations (Pool, 2015). We extended our search to 172 French genomes in PopFly, finding that seven of the DGRP SNPs were absent. Most SNPs, except for those in *gem3, ash2, CG33136, Ddr, CCKLR-17D3* and *lar*, segregated at <0.1. All sites had missing data (>17%, < 96%) across the French genomes. Conversely, 34 out of 59 DGRP SNPs in the Zambian population had no missing data, and nine SNPs had less than 10% missing data. We also examined whether DNA near the SNPs showed adaptive functional differences between the DGRP and other populations using the *F*_ST_ (fixation index) estimator embedded in PopFly. We estimated *F*_ST_ values for the one kb sequence surrounding SNPs with a minor allele frequency higher than 5% in DGRP and Zambian populations. The Ethiopian population was included as another ancestral African population. Mean pairwise comparisons yielded *F*_ST_ values of 0.154 ± 0.0153, and 0.179 ± 0.0292, respectively between DGRP and Zambian or Ethiopian populations, respectively. There was approximately 3-fold less differentiation between the American and French populations (*F*_ST_ = 0.0461 ± 0.01304, z = 4.67399, p < 0.00001, Mann-Whitney U test), showing that despite greater conservation of resistance associated SNPs with ancestral African populations, the DGRP shows less population differentiation with French than ancestral African populations in the surrounding DNA regions.

This is surprising since North American populations contain African and European DNA, but it aligns with comparative studies on microsatellites that showed mean pairwise comparisons between African and non-African populations produce high *F*_ST_ values ranging from 0.11 to 0.20, while European and American populations have average pairwise *F*_ST_ values of 0.071 (Caracristi & Schlötterer, 2003).

## Discussion

Our study broadens previous research on disease resistance variation in the DGRP from North Carolina, United States, to global populations, reflecting *D. melanogaster* evolution as a human commensal. We used 43 global stocks established at different times, acclimated in the laboratory and assayed under standard laboratory conditions, and found persistent latitudinal clines for disease resistance based on their original collection sites. A caveat applying to this research is that lines are likely to suffer from laboratory adaptation and inbreeding. This may not be an issue as previous studies have found that laboratory-maintained inbred *Drosophila* populations exhibit similar trait correlations and plasticity in morphological traits and stresses as freshly caught populations (Maclean et al., 2018), which is as expected for clinal traits shaped by natural selection (Souto-Maior et al., 2020), and we found no evidence of a relationship between stock age and disease resistance in our data. We also took the precaution of testing multiple stocks to provide sufficient independent data points. Finally, as noted by other authors using laboratory stocks to study clinal signals (Fabian et al., 2015), if laboratory adaptation is present at all, it should act against (or even erase) the clinal signals we have detected here.

Our data indicate that disease resistance is linked to collection sites with proximity to the equator and with higher average annual temperatures, as described previously for sleep (Brown et al., 2018). To the 22 strains used by Brown et al (Brown et al., 2018), we added 21 lines, including African populations to provide representation from near the ancestral range of the species (Accra, Ghana). These Ghana lines cluster with Zimbabwe and Zambia populations and contributed to the discovery that sub-Saharan populations are more diverse than other populations (Lack et al., 2016; Pool, 2015; Verspoor & Haddrill, 2011). Accra, due to its equatorial location within the TSGSS biome, experiences minimal seasonal temperature and humidity fluctuations (96% of the year falling within the preferred 20-30°C temperature and 60-100% humidity range), closely resembling the stable, aseasonal Monkey Hill environment. An alternative to latitudinal clines in disease resistance is the possibility that distance (in any direction) away from the range center is associated with altered species interactions and reduced genetic diversity, which may translate to reduced disease resistance. Under this “range center” distance from Africa should negatively correlate with disease resistance to a similar extent as latitudinal clines, which it does not even though latitude is a component of distance. The tendency of lower latitude flies to exhibit higher resistance persisted over large distances suggesting that latitude or climatic variables are more important at predicting disease resistance than a population’s position relative to the species range.

Given the African origin of *D. melanogaster*, the non-African population diversity is expected to be a subset of African diversity (Sprengelmeyer et al., 2019). However, 12(13) of the 15 African male (female) lines were among the 50% most resistant. The exceptions were flies from Madagascar and Zimbabwe and male flies from Malawi, which experience greater seasonal fluctuations in temperature and especially humidity. This indicates that African genomic diversity did not translate into phenotypic diversity in resistance to Ma549 as they were generally resistant, perhaps reflecting their aseasonal habitat. Similarly, four African *D. melanogaster* lines were more resistant to the fungal pathogen *Beauveria bassiana* than two non-African populations (Tinsley et al., 2006). The authors noted that if such variation in pathogen susceptibility also exists in pest insects, it could lead to the evolution of resistance to fungi developed as biopesticides (Tinsley et al., 2006). The frequency of SNPs associated with resistance to Ma549 in the DGRP was lower than that in African genomes, despite nearby DNA resembling European populations. This could be due to African SNPs being favored in the DGRP or even *de novo* mutations in America, as most were rare or absent in France.

African populations, as well as other aseasonal (tropical) populations from uniform climates, such as MH, are generally more disease resistant than seasonal (temperate) populations that endure intermittent thermal and humidity stress. Relatively little is known about clines whose evolution may be governed by interspecific interactions. However, the differences we found between *Drosophila* populations align with studies linking climate variables to malaria risk; uniform temperatures and high humidity support *Anopheles gambiae* vector populations and influence malaria transmission patterns (Paaijmans et al., 2009), emphasizing the importance of aseasonal regions in disease dynamics. Moreover, extensive clinal variation exists within *A. gambiae* populations along an aridity gradient (Cheng et al., 2012). *D. melanogaster* thrives within an optimal temperature range crucial for activity and development (Hamada et al., 2008; Sayeed & Benzer, 1996), while fluctuating temperatures can shorten the lifespan of *D. melanogaster* (Siddiqui & Barlow, 1972). Relative humidity above 60% also significantly contributes to fly longevity, particularly in females (Tochen et al., 2016). Several studies have found that *Drosophila* fly populations increase with rainfall (Achumi et al., 2013; Bombin & Reed, 2016), and fungal insect diseases, including Ma549, are strongly favored by humid conditions (Wang et al., 2023), suggesting a connection between precipitation and increased disease exposure. Aseasonal regions with stable climates and abundant food resources host a majority of terrestrial biodiversity and species interactions (Brown, 2014; Salazar-Mendoza et al., 2021), and their greater species richness and microbial prevalence may select for enhanced host defenses (Schemske et al., 2009). This ecological context of a simultaneous increase of fungal infestations and conditions favorable to fungal infections may act as a stronger selective force at lower latitudes. Similarly, genetically based plant defenses against herbivory are higher at lower latitudes corresponding to greater levels of herbivory in the tropics (Pennings et al., 2009).

Resistance to abiotic stress is likely to be a larger component of fitness at high latitudes. Stress responses are costly, and surviving abiotic stresses requires nutrients and may even reduce immune function (Landis et al., 2012). Thus, alternatively, or in addition, while adapting to abiotic stresses is crucial for the persistence of temperate populations, these adaptive processes could be traded-off against alleles imparting greater resistance to disease. Hoffmann et al. have proposed a similar evolutionary trade-off where an individual fly cannot be both cold resistant and starvation resistant, with starvation resistance being selected for in warm climates (Hoffmann et al., 2005). Our previous DGRP analysis found an association between starvation resistance and disease resistance (Wang et al., 2017), suggesting that it could be one of the factors contributing to adaptive differences in disease resistance. Starvation resistance is linked with metabolic storage and the ability to maintain homeostasis, which could help deal with abiotic and biotic stressors. However, high resource availability and pathogen prevalence in tropical climates could also select for disease resistance. Another study of the DGRPs found significant correlations between abdomen size at 28°C and chill coma recovery and between abdomen size at 17°C and resistance to Ma549 (Lafuente et al., 2018), suggesting that disease resistance is linked to developmental plasticity that helps flies cope with environmental heterogeneity. Most of the variation in the DGRP that predicted resistance affected the general robustness of the flies, and in particular genes involved in development, nutrition and sleep indices (Wang et al., 2017).

Females are more dependent on relative humidity values above 60% than males (Tochen et al., 2016). In this study, we used previously acquired desiccation and disease resistance data from DGRP lines (Rajpurohit et al., 2018; Wang et al., 2017) to demonstrate a significant link between them in females, indicative of cross-resistance to multiple stresses rather than tradeoffs, and therefore common mechanisms for some of the variation in these traits. The resistance of DGRP lines to the bacterium *Pseudomonas aeruginosa* and the narrow host range *Entomophthora muscae* (Entomophthoromycota) are correlated with resistance to Ma549, indicating the presence of general (non-specific multipurpose) defenses and that resistance to one pathogen is not traded off with increased susceptibility to a different pathogen (Wang et al., 2020; Wang et al., 2017).

We have assumed that disease resistance is linked to adaptation to local environments. Local adaptation seems to be the case for desiccation resistance as laboratory conditions that mimic the low humidity of IP favor IP flies over MH flies. However, if some insects have evolved an increased generalized resistance to multiple stressors, then resistance to Ma549 may be a co-incidental side effect of cross-resistance to other stressors and not derived from an evolutionary history with pathogens. The fly immune system is known to have functions beyond defense against pathogens with cold stress in particular known to activate immune genes and increase resistance to *B. bassiana* (Wiil et al., 2023). Likewise, reactive oxygen species activate some innate immune mechanisms (Ramond et al., 2021), as well as sleep, which protects against oxidative stress (Hill et al., 2018), so not surprisingly, the resistance of DGRP lines to oxidative stress is correlated with resistance to Ma549 (Wang et al., 2017).

It is unknown whether this multi-stress resistance is still advantageous when insects simultaneously encounter abiotic and biotic stresses. This scenario is most likely to occur in seasonal regions due to arid or thermal stresses. Although desiccation and disease resistance are associated in females, disease resistance peaks in stable regions where desiccation is not a major threat. Flies from IP and males from MH represent the range of variation in disease resistance. Monkey Hill’s stable, warm, near-optimal temperature and humidity (marine ecoregion) presumably imposes less abiotic stress than the seasonally variable, drier desert climate of IP (DXS biome), while pathogen pressure is likely to be low in an arid desert. Relaxation of selective pressure could have led to degeneration of immune defenses in IP flies, similar to *D. sechellia* that lives in an “enemy-free space” (O’Malley et al., 2023). Conversely, IP flies display high desiccation resistance with the highly disease-resistant MH males being the most susceptible to desiccation (Hoffmann & Harshman, 1999). This reversal in the direction of sexual dimorphism aligns with the DGRP lines (Figure 5E-F) and a recent study on sexual dimorphism in 15 *Drosophila* lines, which found that males generally live longer but females are more resistant to starvation and desiccation, with greater variation between lines in these phenotypes (Lin et al., 2023). These findings indicate that the relationship between desiccation and disease resistance is prioritized differently in males and females.

We found that the association between African populations and DGRP was particularly strong for SNPs linked to the phenotypic micro-environmental plasticity of DGRP lines. DGRP lines exhibit significant variability in their plasticity to disease (Wang et al., 2017) and thermal stress (Lafuente et al., 2018), showing that even within a local population, genotypes can produce flies with different plasticity. Plasticity is generally costly and only selected for in heterogenous environments (van den Heuvel et al., 2013), which implies males, that usually showed greater variation in disease resistance, may be more adaptable or experience a greater diversity of pathogens. Using *CV_E_*, we found that selection favors the plasticity of disease resistance in males equally in seasonal and aseasonal populations, whereas in females, there was a trend for reduced plasticity in stable climates. This implies that aseasonal females may be less resilient to changes in disease exposure. Female fitness is more critical for population persistence and growth than male fitness (Harts et al., 2014), so these differences may affect species survival when confronted with global climate change and infectious diseases (St. Leger, 2021).

A central problem in infection biology is determining why two individuals exposed to the same pathogen have different outcomes. We found that the resistant MH, the highly susceptible IP, and a line (CG), which shows almost no sexual dimorphism in disease resistance, had marked differences in their ability to control Ma549 growth and colonization rate, and thus pathogen invasiveness. Because variation in survival is linked to fly robustness and resistance to stressors, as well as physiology and behaviors such as sleep, we had expected flies to differ in their ability to tolerate extensive colonization. Instead, all lines showed low fungal loads until the day before death, suggesting that they die at a specific fungal load. Thus, variation in survival stems from variations in the ability to repress fungal growth, which is most likely determined by the immune system rather than the ability to tolerate extensive colonization. The anti-microbial peptides (AMPs) Bomanins only found in *Drosophila* spp are candidates for this role as unlike other *Drosophila* AMPs they have activity against Ma549 (Wang et al., 2023). Classical virulence theory suggests that the extra resistance demonstrated by MH could drive selection for higher pathogen virulence, while tolerance, if it had occurred, could have led to eventual pathogen-host co-existence (Howick & Lazzaro, 2014).

Using sleep data for healthy females from 22 lines (Brown et al., 2018), we demonstrated that variation in disease resistance was positively correlated with consolidated sleep (sleep duration and bout length) and negatively correlated with fragmented sleep (bout number). This suggests that flies that sleep more while healthy are better at resisting infection. In addition to the diurnal rhythm sleep observed in healthy animals, *Drosophila* increases sleep in response to infection, akin to rabbits reacting to fungal infection (Toth et al., 1993). This “sickness sleep” precedes symptoms like reduced feeding and the upregulation of the immune gene *Drosomycin* during Ma549 infection (Lu et al., 2015; Wang et al., 2023). Our sleep analysis showed Ma549 infection increased daytime sickness sleep in MH, CG, and IP flies, especially in males, most notably MH males who succumbed the slowest to Ma549. The benefits of sleep for diseased organisms are unclear, but the GWAS using the DGRP found overlapping alleles affecting resistance to Ma549 and sleep, some of which are sex-dependent, impacting sleep and/or disease resistance in only one sex (Wang et al., 2017). The differences between male and female MH, CG, and IP flies may be a phenotypic expression of this phenomenon. Males exhibited a lower fungal burden than females, consistent with sex-specific immune investment strategies. Sexual activity reduced sleep and the ability to delay fungal growth in both sexes, consistent with trade-offs between reproduction and immunity (Garbe et al., 2016; Gordon et al., 2022; Rai et al., 2023), even though males do not experience the energy drain of egg production (Mishra et al., 2023).

The diverse mechanisms involved in disease resistance have typically been studied independently, limiting our understanding of the interplay between biotic and abiotic stress responses and their relation to disease resistance and ecology. The issue is particularly relevant for clines in sleep, stress resistance, and defense, as these traits are closely linked to fitness. Growing recognition exists that stress and defense traits are interconnected with each other, and with sleep, yet the general issue of their joint clinal evolution remains unresolved. Given the anticipated climate change and the importance of rapid evolution in invasive species, this is likely to change, and the *Drosophila*-Ma549 model appears to be well-suited for these studies. Sleep and stress responses may provide an evolutionary mechanism for optimizing host defenses, but it is unclear whether selection primarily shapes host defenses through changes in non-specific stress responses and variations in physiological and behavioral traits such as sleep or vice versa whether the selection is on stress responses or sleep with disease resistance as a secondary factor. IP females exhibit greater desiccation resistance than males, but the high nocturnal activity (low sleep) of IP males might feasibly be a behavioral adaptation to their arid environmental conditions and unrelated to disease resistance. Our findings emphasize the critical role of environmental conditions on fly lifespan post-infection, highlighting the complex relationship between ecological context and disease resistance.

## Materials and Methods

### Fly stocks, geographic and climate data, and biomes

Twenty-two fly populations were a gift from Dr. Alex Keene’s lab (Texas A&M University) (Brown et al., 2018) and an additional 21 fly populations [Ghana (Mataheko or Dansoman, Accra), Mauritius (Le Reduit), Malawi (Lujeri), Madagascar (Tananarive), France (Montpellier), and the United Kingdom (Sussex)]) were acquired from the *Drosophila* Species Stock Center (Cornell University, Ithaca, NY) with all stock numbers given in Table 1. *Drosophila* were cultured on cornmeal-molasses media supplemented with yeast, agar, Tegosept, and propionic acid at 25°C and ∼70% humidity under a 12-hour light:dark cycle. Virgin flies of each genotype were collected under light CO_2_ anesthesia within 8 h of eclosion and housed in separate vials for 2–3 days to mature.

Latitude, longitude, and altitude measurements of each collection site in Table 1 were obtained from Google Earth Pro and converted from degrees, minutes, seconds to decimal degrees. Latitude and altitude measurements were transformed into absolute values, followed by conversion to log_10_ for regression analysis. The 43 fly populations were categorized into six biomes based on the latitude and longitude of each collection site using ArcGIS Pro (2022, Version 3.0.1, Esri Inc., Redlands, CA, USA), and 14 biomes defined in Ecoregions 2017 (Dinerstein et al., 2017). Because latitude, altitude, and precipitation measurements were not normally distributed, we calculated the log_10_ values for the Ma549 LT_50_ regression analyses. The geographic data in Figure 1A were generated using ArcGIS Pro.

Monthly average, high (daytime), and low (nighttime) temperature datasets of land and ocean in a 1° × 1° latitude-longitude grid covering the Earth’s surface from 1850 to the present were obtained from the Berkeley Earth website (http://berkeleyearth.org/data/) and verified using the visualization tool L3HARRIS IDL software (https://www.l3harrisgeospatial.com/Software-Technology/IDL). Temperature data based on collection year of all 28 collection sites were in a gridded NetCDF format and extracted with a self-coded program script (S2B Table), as provided in S2A Table. The annual average, high, and low temperatures were computed by summing the respective monthly averages, highs, and lows and dividing each sum by 12.

The monthly precipitation data within a 0.5° grid for each collection site and collection year in the S2A Table were extracted from a large database of up to 10 different weather stations at the World Meteorological Organization (Schneider et al., 2020). The annual mean precipitation was determined by averaging the monthly data across all 12 months. The month with the highest precipitation value among the 12 months was identified as the wettest month.

Data on 52,584 hourly temperatures and relative humidity’s from the years 2005-2010 for all 28 collection sites defined by latitude and longitude were obtained from the Prediction of Worldwide Energy Resource (POWER) Project, Langley Research Center, National Aeronautics and Space Administration. To cover all 43 fly populations, we used NASA POWER data to analyze hourly temperature and humidity from 2005 to 2010 at 28 collection sites, aligning with the collection dates of most fly lines (S3 Table). This data was utilized to analyze the effects of environmental factors on fly longevity. The total hours during which temperatures fell within specific ranges and relative humidities within 60-100% from 2005-2010 across all 28 collection sites were calculated using 52,584 hourly temperature and humidity data points. The total hours were then divided by six years to calculate the annual hourly duration (hours/year) for 43 fly lines, which were used in simple linear regression analyses.

### Fungal strains, and longevity bioassays

*Metarhizium anisopliae* 549 (Ma549) was acquired from the ARS Collection of Entomopathogenic Fungal Cultures (ARSEF) (Ithaca, NY). Ma549 cultures were thawed from −80°C stock vials and grown for 10 to 14 days at 27 °C on potato-dextrose-agar media plates.

Bioassays using *M. anisopliae* 549 transformed to express green fluorescent protein (Ma549-GFP) were performed according to a previous report (Lu et al., 2015) on 43 worldwide fly populations. Briefly, fly populations were infected at the same time of day and survival was monitored after topical inoculation in groups of five replicates (20-30 flies each) per sex per line. Each line was screened three times. To prepare inoculum, conidia were suspended in sterile distilled water, vortexed for 2 minutes and filtered through Miracloth (22-25µm) (Andwin Scientific). Flies (two to four days old) were vortexed with spore suspensions (20 ml, 2.5×10^4^ conidia/ml) for 30 seconds, collected by filtering through Miracloth, and transferred into vials containing fly stock food without Tegosept and propionic acid. Flies were cultured at 27°C and ∼85% relative humidity. Male and female flies were housed in separate food vials to examine sexual dimorphism and avoid an interaction between the effects of sex and age in response to infection. Less than 10% of flies vortexed with water alone (mock-infected), or conidial suspensions died within one day, and there were no significant differences between lines; therefore, flies succumbing within one day post infection were deleted from the infection data. The flies were flipped into fresh food vials daily. The number of dead flies was recorded twice daily for 10-14 days. Host survival differences were measured as LT_50_ values (median lethal time in days at which 50% of the flies died) calculated using R 4.1.0 (R Core Team 2021). This method was highly reproducible, with a mean LT_50_ value for control *Drs*-GFP flies of 4.682 ± 0.029 (N = 126). The inoculum load of spores per male and female *Drosophila* was approximately 200 colony-forming units (Lu et al., 2015). To quantify micro-environmental plasticity in mean survival time (MST) as described in a previous report (Morgante et al., 2015), we used either the untransformed or transformed within-line standard deviation (*σ_E_*) and coefficient of environmental variation (*CV_E_*) (S1B Table), depending on normality tests. *σ_E_* has the advantage of being measured in days post-infection, the same scale as the trait mean, while *CV_E_* is calculated as the standard deviation divided by the mean and multiplied by 100.

### Desiccation assays

Ten age-matched (three to six days old) flies were sorted by sex using light CO_2_ anesthesia and placed into a 25 x 95 mm tube, covered at the top with gauze, and joined with parafilm to another 25 x 95 mm tube containing 10 g of silica gel desiccant beads (Wisedry B01MPYB16J) at the bottom to achieve a desiccated relative humidity of 5-10%. Two replicates per sex per line were used for each experimental trial. Control chambers contained 5 ml of a supersaturated NaCl solution to maintain a stable relative humidity of approximately 70%. The flies were monitored at one-hour intervals for 20 hours for death, as indicated by failure to right themselves or to move their legs when their vials were tapped or inverted (Gibbs et al., 1997).

### Fungal colonization of hosts measured by colony forming units (CFUs)

A time-course bioassay of fungal growth in the hemolymph of mated and virgin MH, CG, and IP females and males was conducted using previously described protocols (Lu et al., 2015). Fifteen flies per sex, mating status, and line at each time point across three experiments were individually homogenized with 45 μl of 0.1% Tween 80. For the resistant MH line, the entire 45 μl homogenate was spread onto *Metarhizium* selective medium (Rose Bengal Agar plates supplemented with oxbile, CTAB, oxytetracycline, streptomycin, penicillin, chloramphenicol, and cycloheximide (Hu & St Leger, 2002). For susceptible lines, 5 μl of a 10-fold dilution of the homogenate was spread onto the plates. Colony forming units (CFUs) were counted using the ImageJ Cell Counter after up to 7-days incubation at 25°C.

### Sleep assays and data analysis

The *Drosophila* Activity Monitor (DAM2) from Trikinetics, Waltham, MA, was used to monitor locomotor activity and analyze sleep patterns in MH (resistant), IP (susceptible), and CG (average longevity) flies. Flies were individually housed in 5 x 65 mm monitor tubes containing 5% sucrose and 2% agar after CO_2_ anesthesia, with any initial mortalities replaced within 12 hours to minimize acclimation effects. Monitoring occurred over seven days under a 12-hour light/dark (LD) cycle (lights on at 9:00 AM EDT, lights off at 9:00 PM EDT). Sleep data, defined as periods of 5 minutes or more of inactivity (Shaw et al., 2000), were processed using DAMFileScan113 software (Trikinetics) and analyzed using custom software (Insomniac3, RP Metrix, Skillman, NJ; gift of Dr. Julie Williams, University of Pennsylvania). The DAM2 output was converted to total daily, nighttime, and daytime sleep duration, represented by the number of minutes when a fly was asleep in 24 or 12 hours. Sleep patterns were examined from 12 to 60 hours post-infection with Ma549, spanning two light-dark cycles. This period precedes the onset of most visible symptoms, such as reduced feeding and reproductive behavior, and the upregulation of the immune gene *Drosomycin* in response to Ma549 infection (Lu et al., 2015; Wang et al., 2023). The sleep profiles of mated CG and IP flies were compared with those of MH flies under normal conditions, and infected flies were compared with uninfected peers to assess the impact of infection within each line. Sex-specific differences in sleep and disease outcomes were also investigated along with the influence of mating status on disease resistance and sleep patterns. Sleep differences were quantified using the net percentage differences between the experimental groups (S5 Table).

### Statistical analyses on interactions among clinal cues, geographic variables, longevity, and sleep parameters

We conducted simple linear regression analyses between LT_50_ values and environmental cues or geographic variables (altitude and latitude) at collection sites for 43 global (subdivided into 23 aseasonal and 20 seasonal) populations. We also correlated LT_50_ values with sleep parameters data for 22 female fly populations (Brown et al., 2018). Pearson’s correlation analyses were performed in GraphPad Prism 10 (GraphPad Software, Boston, MA, USA).

To assess the survival rates of flies following Ma549 infection, we conducted Kaplan-Meier survival analysis. Survival curves for each group (MH, CG, and IP), including variations by sex (male or female) and mating status (mated or virgin), were generated using GraphPad Prism 10. Statistical significance between survival curves was determined using the log-rank (Mantel-Cox) test and Gehan-Breslow-Wilcoxon test. Percentage survival and standard errors were calculated and compared across different genotypes and conditions to assess the impact of genetic variation and infection on fly survival.

We evaluated data normality using the D’Agostino-Pearson and Shapiro–Wilk normality tests, known for their robustness across various statistical distributions (Yap & Sim, 2011). For data sets passing at least one of normality test and showing statistically similar standard deviations (SDs), we employed the unpaired Student’s t-test for two group comparisons and one-way ANOVA (Tukey’s multiple comparisons) for three or more groups. If variances significantly differed (p < 0.05), we used Welch’s correction (Welch’s t-test) for two groups and the one-way Welch’s ANOVA (Dunnett’s multiple comparisons) for multiple groups. For data sets failing normality tests, we used the Mann–Whitney U test for two groups and the Kruskal-Wallis test with Dunn’s post hoc test for three or more groups. Reported values included p-values, median ± IQR (interquartile range: the difference between the 75th and 25th percentile) from the Kruskal-Wallis test with Dunn’s post hoc multiple comparisons test, and mean ± SEM from ordinary ANOVA or Welch’s test. Statistical significance was set at p < 0.05.

JMP^®^ Pro, version 17.1 (SAS Institute Inc., Cary, NC, USA) was utilized for the forward stepwise regression analysis on the impact of geography and biome on resistance levels (Table 3), and principal component loadings of environmental variables and LT_50_ for male and female flies (Table 4), as well as ANOVA involving four factors and generating interaction profiles (Appendix 4—table 1 and Figure 8—figure supplement 1).

## Acknowledgments

We would like to thank Dr. Kelly A. Hamby (University of Maryland) for her valuable input on biome concepts and the insightful discussions, Drs. Elizabeth Brown and Alex Keene (Texas A&M University) for providing 22 worldwide fly stocks, Dr. Julie Williams (University of Pennsylvania) for providing insomniac sleep analysis software, Mr. Yufeng Zhu (National Oceanic and Atmospheric Administration) for his help on extracting and verifying large temperature and precipitation datasets from the 28 collection sites.

## Author Contributions

**Conceptualization**: Mintong Nan, Raymond J. St. Leger.

**Data curation**: Mintong Nan, Jonathan B. Wang, Raymond J. St. Leger.

**Formal analysis**: Mintong Nan, Raymond J. St. Leger.

**Funding acquisition**: Raymond J. St. Leger.

**Investigation**: Mintong Nan, Jonathan B. Wang.

**Methodology**: Mintong Nan.

**Project administration**: Raymond J. St. Leger.

**Resources**: Mintong Nan, Jonathan B. Wang, Raymond J. St. Leger.

**Supervision**: Raymond J. St. Leger.

**Validation:** Mintong Nan, Raymond J. St. Leger.

**Visualization**: Mintong Nan.

**Writing** – original draft: Mintong Nan, Raymond J. St. Leger.

**Writing** – review & editing: Mintong Nan, Raymond J. St. Leger.

## Supplementary files

S1 Table. Survival data following Ma549 infection for 43 global mated females and males, as well as virgin CG, IP, and MH females and males.

S2 Table. Monthly average, high, and low temperatures (°C), precipitation and precipitation in wettest month at the collection year of each of 43 lines, along with the ncPick script.

S3 Table. Hourly temperature and relative humidity data from the years 2005-2010 for 28 locales or 43 fly strains.

S4 Table. Desiccation data for IP and MH strains, along with disease and desiccation resistance values in 162 DGRP lines in Figure 5.

S5 Table. Sleep data in Figures 7-8.

S6 Table. CFU data in Figure 9B-D.

S7 Table. LT_50_ and CV_E_ SNP frequency in Figure 10.

S8 Table-ANOVA summaries on main effects and interactions.

---

**Appendix 1—table 1.**
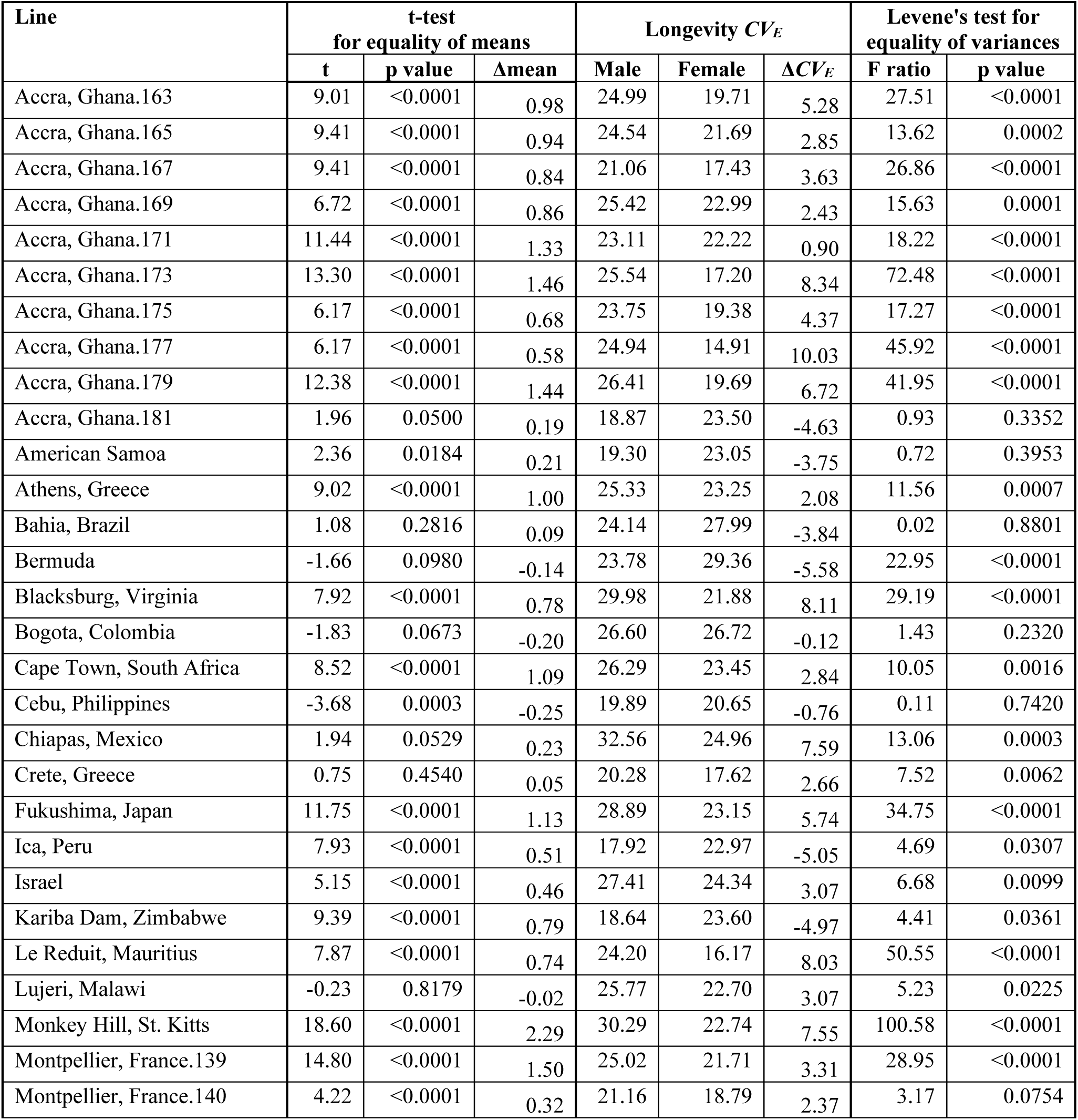
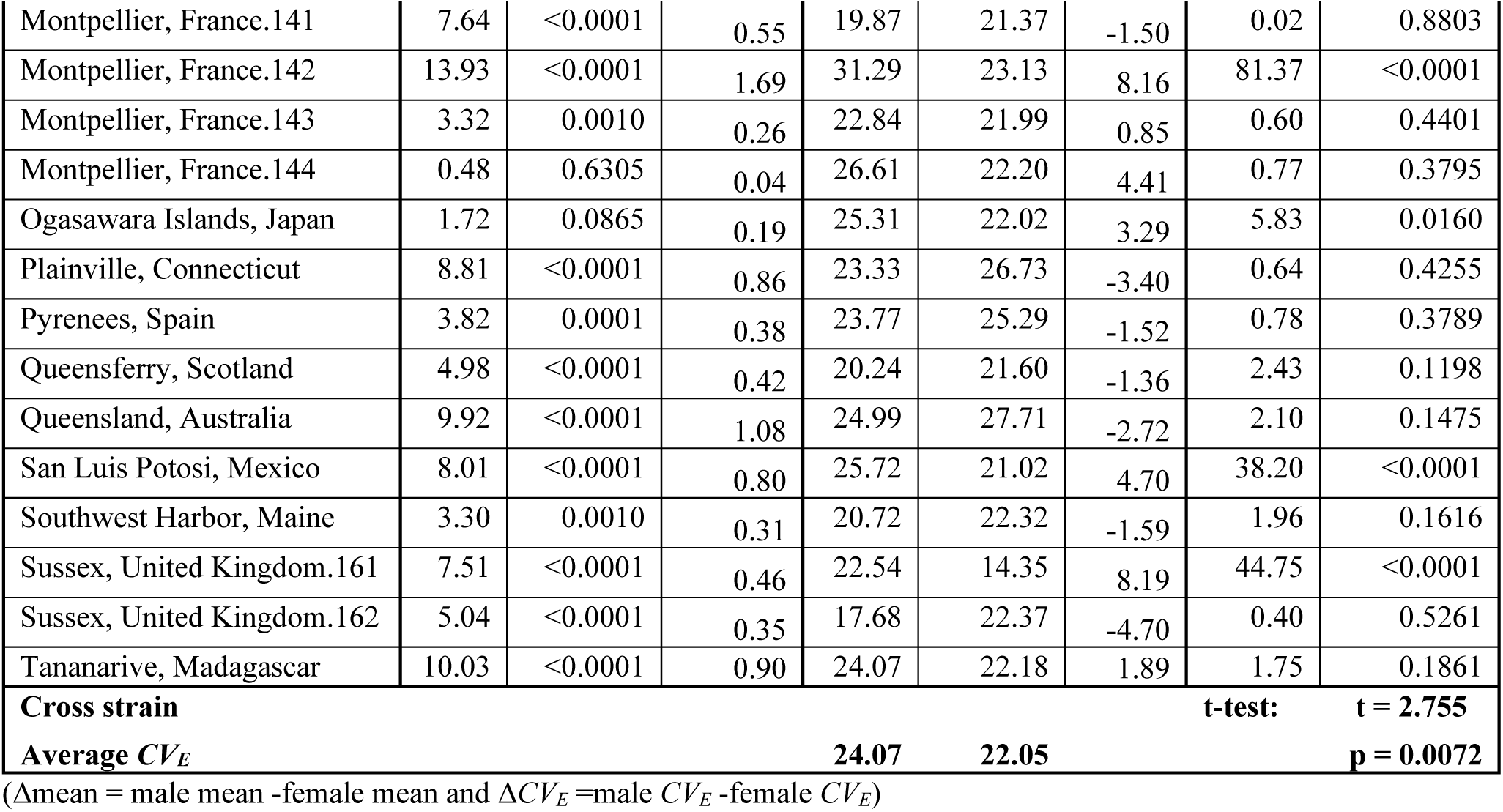
Comparison of mean and variation in disease resistance between males and females.

**Figure 2—figure supplement 1.**
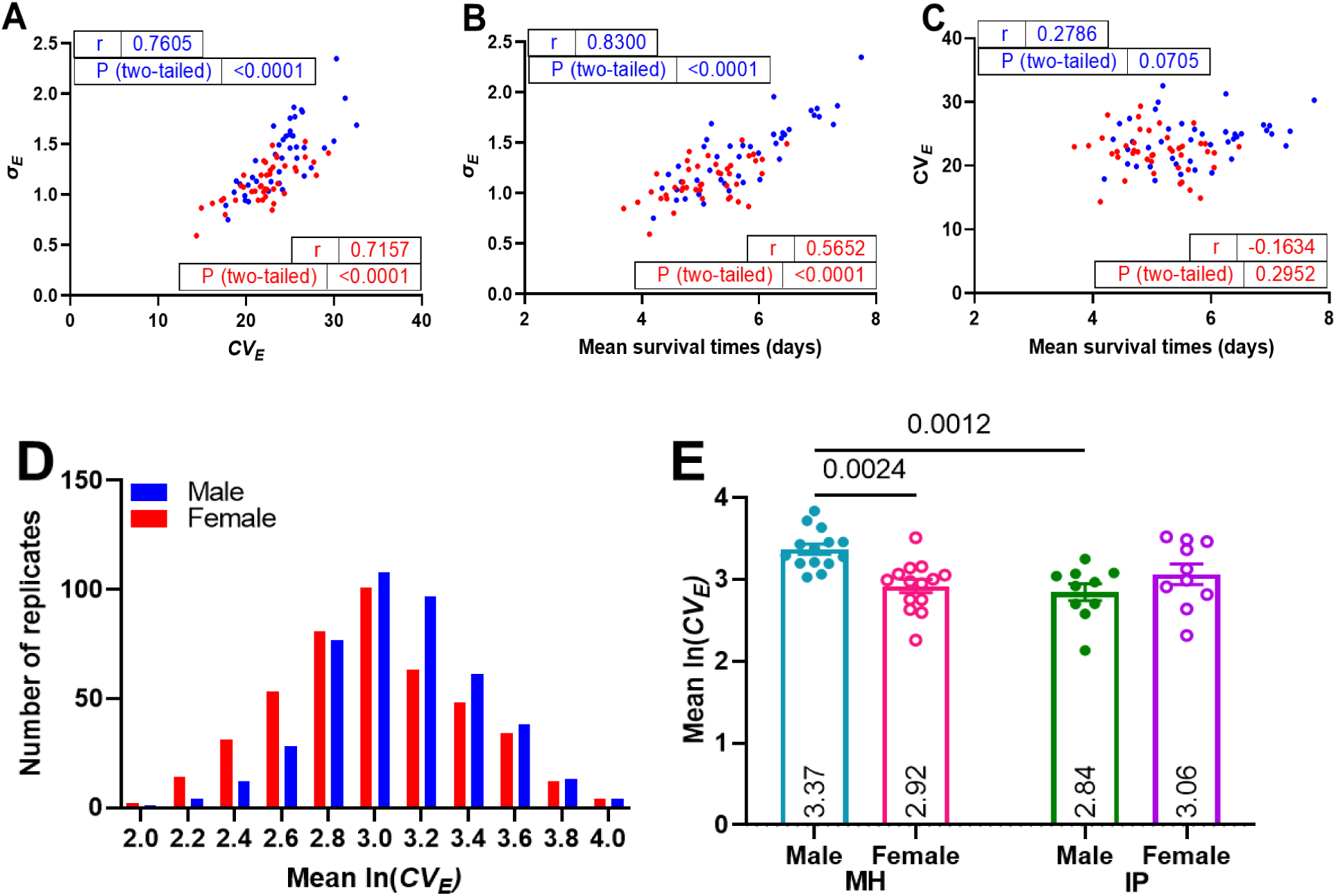
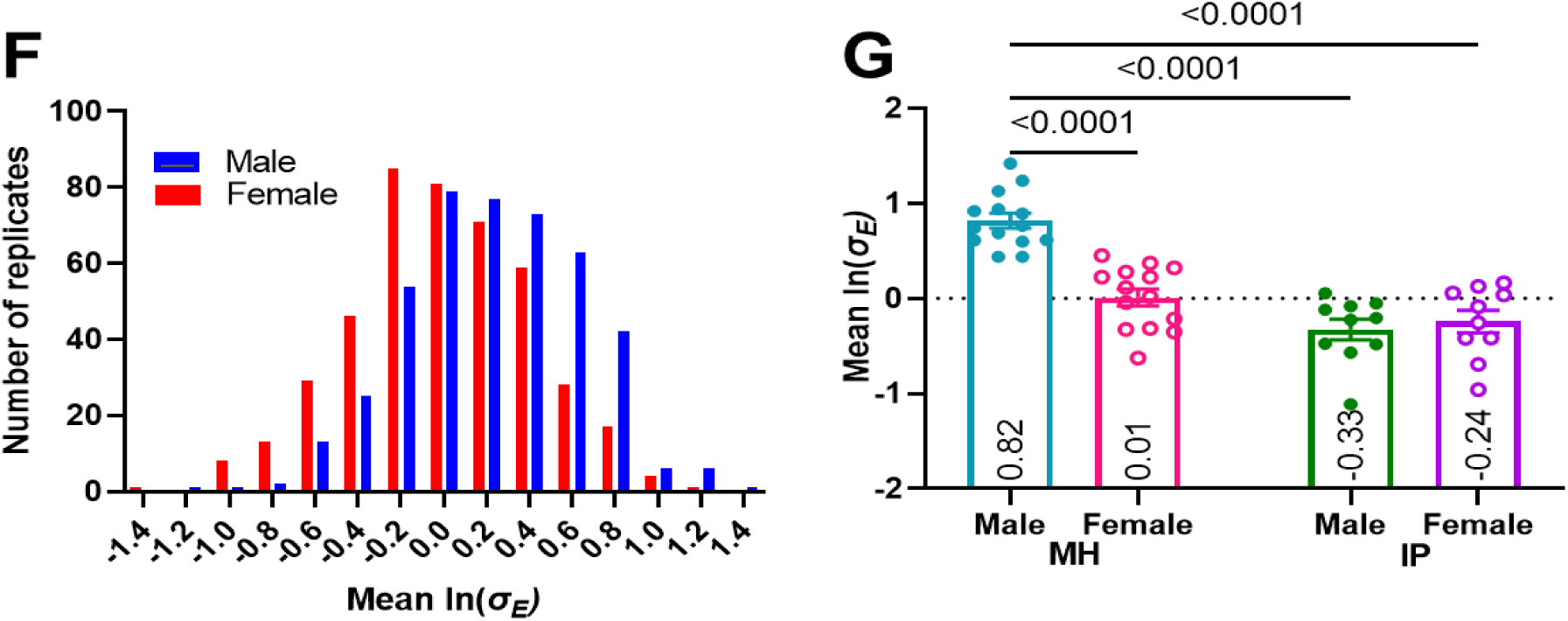
Correlation analyses of plasticity in global lines exposed to Ma549 are presented for male and female *σ_E_* versus *CV_E_* **(A)**, *σ_E_*versus MST **(B)**, and *CV_E_* versus MST **(C)**, with corresponding r and p-values. Blue and red dots in these figures represent males and females from all 43 lines, respectively. Plasticity distribution **(D, F)** and bar graph **(E, G)** of ln(*CV_E_*) **(D, E)** and ln(σ_E_) **(F, G).** 443 global male (blue) and female (red) replicates are represented in panels D and F, while 14 replicates with 373 MH males (turquoise) and 501 females (crimson) from three trials, 10 replicates with 344 IP males (green) and 281 females (purple) from two trials are depicted in panels E and G. The transformed ln(*σ_E_*) and ln(*CV_E_*) show a normal distribution in D and F. Panels E and G use two-way ANOVA (Tukey’s multiple comparisons) to analyze MH and IP plasticity, presenting mean ± SEM and p-values.

## Appendix 2—figure 1 Geographic variables impact host susceptibility to infection

Two-way ANOVA demonstrated significant main effects on fly longevity, with sex accounting for 5.549% (F_(1, 456)_ = 29.37, p < 0.0001), and geographical origins for 8.296% (F_(4, 456)_ = 10.98, p < 0.0001) of the total variation (S8B Table). DGRP males (females) were 1.09 (1.10)-fold more resistant than European/Middle Eastern males (females) (p = 0.0207).

**Appendix 2—figure 1.**
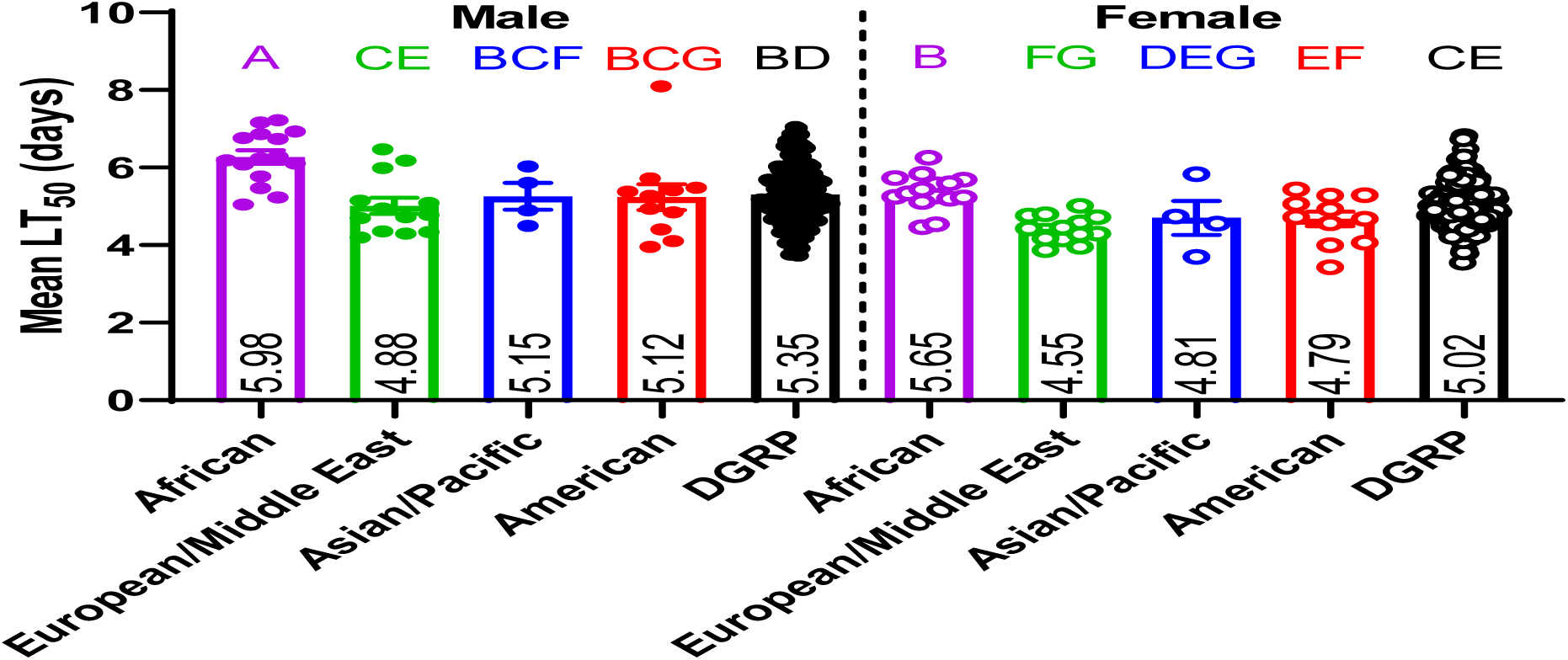
Geographic variation in LT_50_ values among worldwide fly lines infected with Ma549. The 43 worldwide fly lines are depicted with dots: pink for African, green for Europe/Middle East, blue for Asian/Pacific, and red for American regions. LT_50_ values were averaged for each of the 43 male and female lines, with data obtained from 10-15 replicates (averaging 35 flies each) per sex per line across 2-3 experiments. Data from previously published DGRP lines are shown in black (Wang et al., 2017). A two-way ANOVA was conducted to evaluate the main effects of geographic region and sex on disease resistance. The analysis revealed no significant interaction between region and sex (F_(4, 456)_ = 2.261, p = 0.0617). Least squares means (LSM) ± SEM from Tukey’s multiple comparison test are shown, with a compact letter display (CLD) positioned at the top of individual bars, with statistical significance where bars are not connected by the same letter.

## Appendix 3—figure 1 Linear regression analyses of latitude, altitude, and climate variables for global fly collection sites (2005-2010)

Using NASA Power Data, we analyzed hourly temperature and relative humidity from 2005 to 2010 at 28 collection sites, corresponding to the collection dates of most of the 43 fly lines (S3 Table). Using NASA Power Data to analyze hourly temperature, we are able to detect very short interludes with extreme conditions, such as a reading of 43.46°C in Australia. We recorded mean annual hourly minimum, maximum, and range (maximum - minimum) values. Linear regressions show strong correlations between collection site latitude (Table 1) and minimum annual temperature (Tmin) (**A**) and annual temperature range (Trange) (**C**), but not maximum annual temperature (Tmax) (**B**). Altitude (Table 1) moderately correlates with minimum annual relative humidity (RHmin) (**D**) and annual relative humidity range (RHrange) (**F**), but not maximum annual relative humidity (RHmax) (**E**). The narrow ranges of maximum annual temperature and humidity explain the lack of correlations with latitude or altitude.

**Appendix 3—figure 1.**
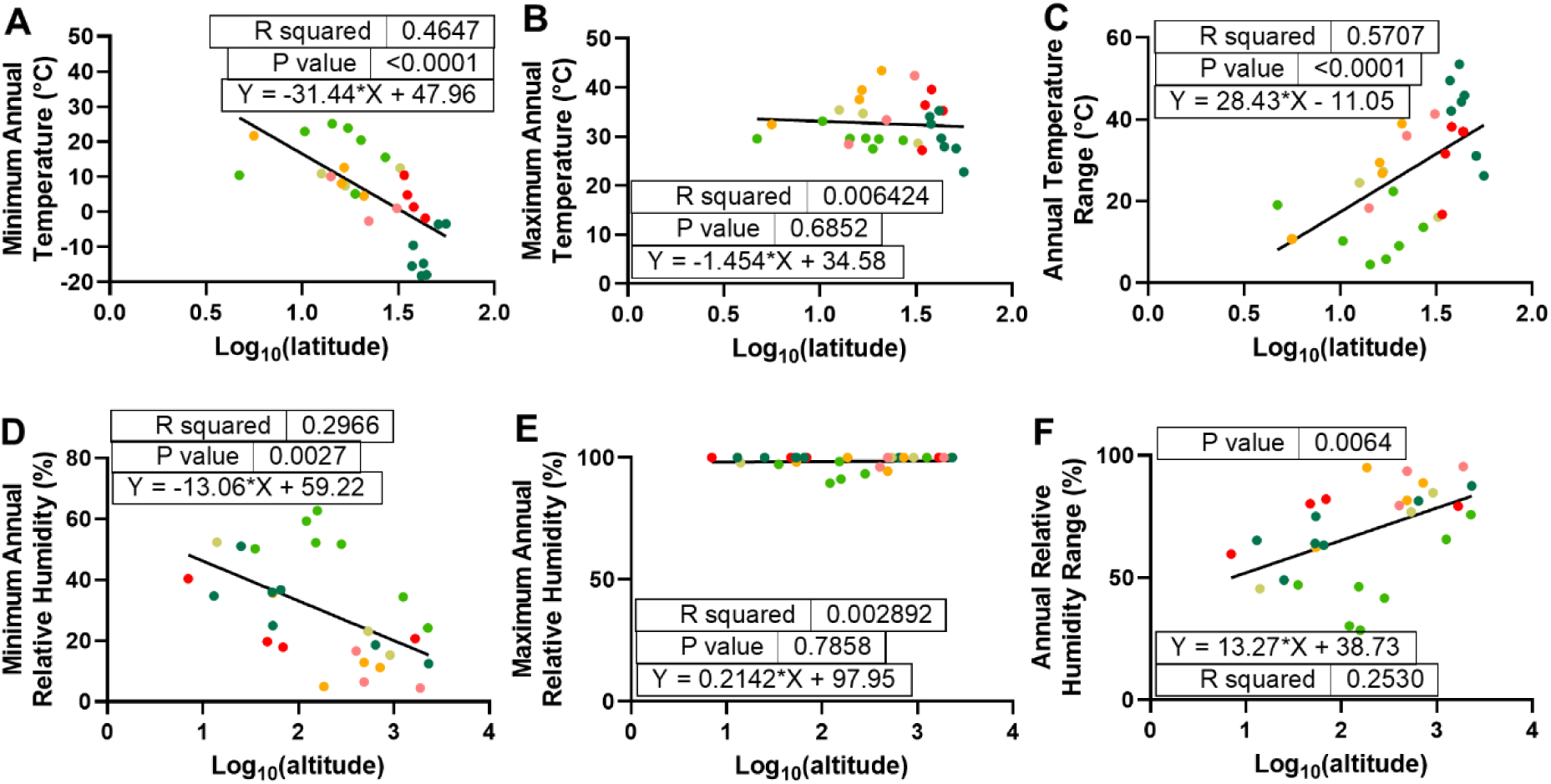
Simple linear regression analyses between latitude and annual temperatures or altitude and relative humidities for the global fly population collection sites: Biome colors across 28 geographic locations (2005-2010) match those in Figure 1A.The figures show results for minimum annual temperature **(A)**, maximum annual temperature **(B)**, annual temperature range **(C)**, minimum annual relative humidity **(D)**, maximum annual relative humidity **(E)**, and annual relative humidity range **(F)**, including R^2^ values, p-values, and regression equations with slopes.

## Appendix 3—figure 2 Simple linear regression analyses correlating Ma549 LT50 values with latitude, altitude, annual temperatures or precipitations at collection sites for global fly populations

**Appendix 3—figure 2.**
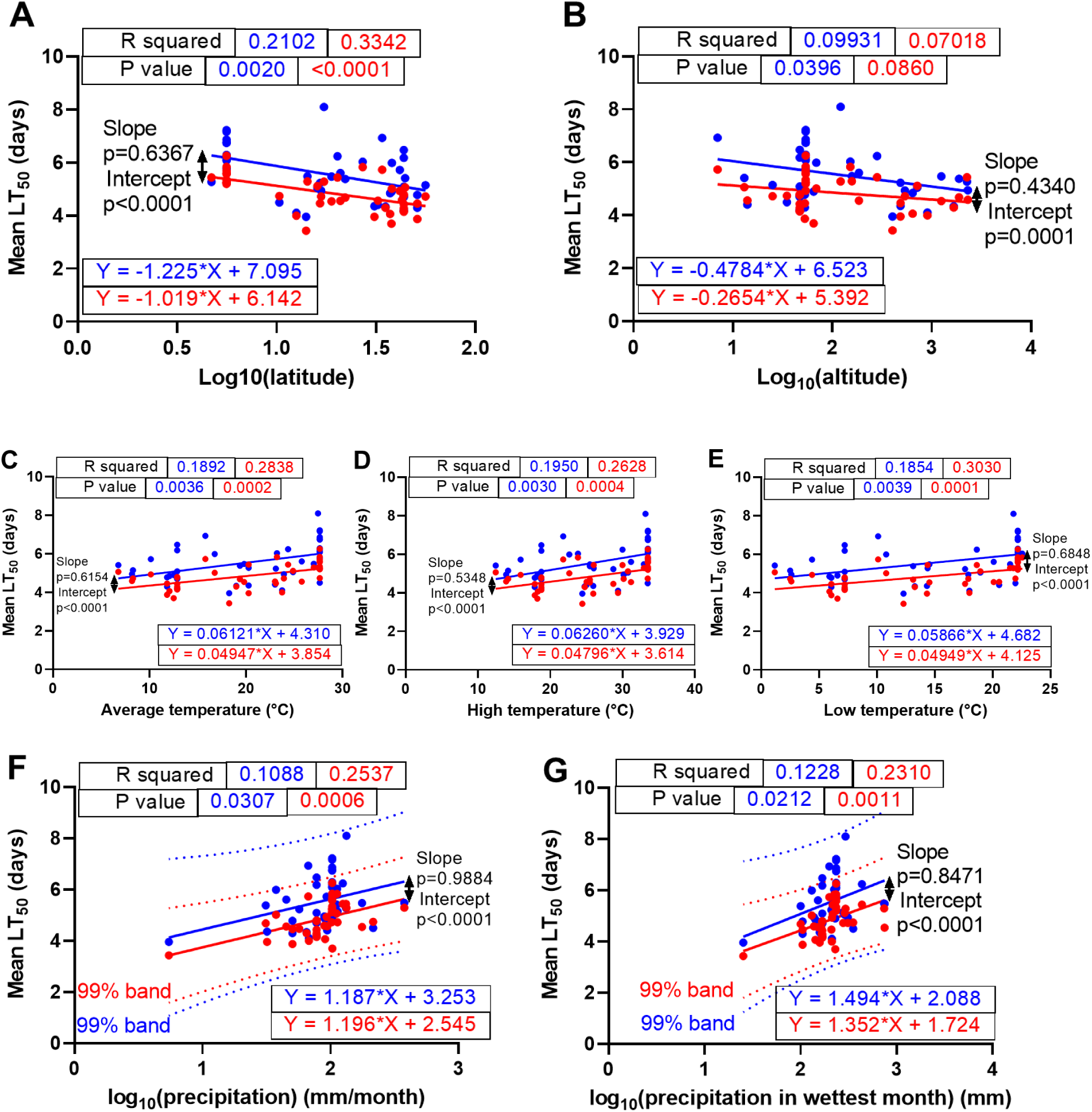
Simple linear regression analyses correlating Ma549 LT_50_ values with latitude, altitude, annual temperatures or precipitations at collection sites for global fly populations: Temperature and precipitation data were extracted from all 28 collection sites based on the original collection year for each fly line. Results for latitude (**A**), altitude (**B**), annual average temperatures (**C**), high temperature (**D**), low temperature (**E**), annual mean precipitation (**F**), and precipitation in wettest month (**G**) are shown, featuring R^2^ values, p-values, and regression equations including slopes. Male (female) regression lines are in blue (red), with p-values for comparing slopes and intercepts between female and male lines using analysis of covariance (ANCOVA).

## Figure 3—figure supplement 1 The effects of annual temperature and relative humidity on female and male LT_50_ and plasticity values

To investigate disease resistance, we calculated the average annual exposure to the preferred temperature (20-30°C) using 52,584 hourly data points from 2005 to 2010 (S3A Table). Exposure at collection sites ranged from 75 to 8,764 hours/year (A-C).

Linear regression analyses for the 43 fly populations showed strong correlations between mean survival time (MST) in response to Ma549 for males and females and experiencing longer aseasonal uniformity measured as the hours per year spent within preferred temperature ranges (**A**). Positive correlations were also found between *σ_E_* and increasing aseasonal uniformity in males, but not females (**B**). We observed no evidence of correlation between *CV_E_* values and increasing aseasonal uniformity in males and females (**C**). Despite differing intercepts due to males’ greater resistance, the MST values for both sexes trended similarly with temperature, resulting in nearly parallel regression slopes. Higher R² values for females suggest that temperature have a greater impact on female disease resistance.

**Figure 3—figure supplement 2.**
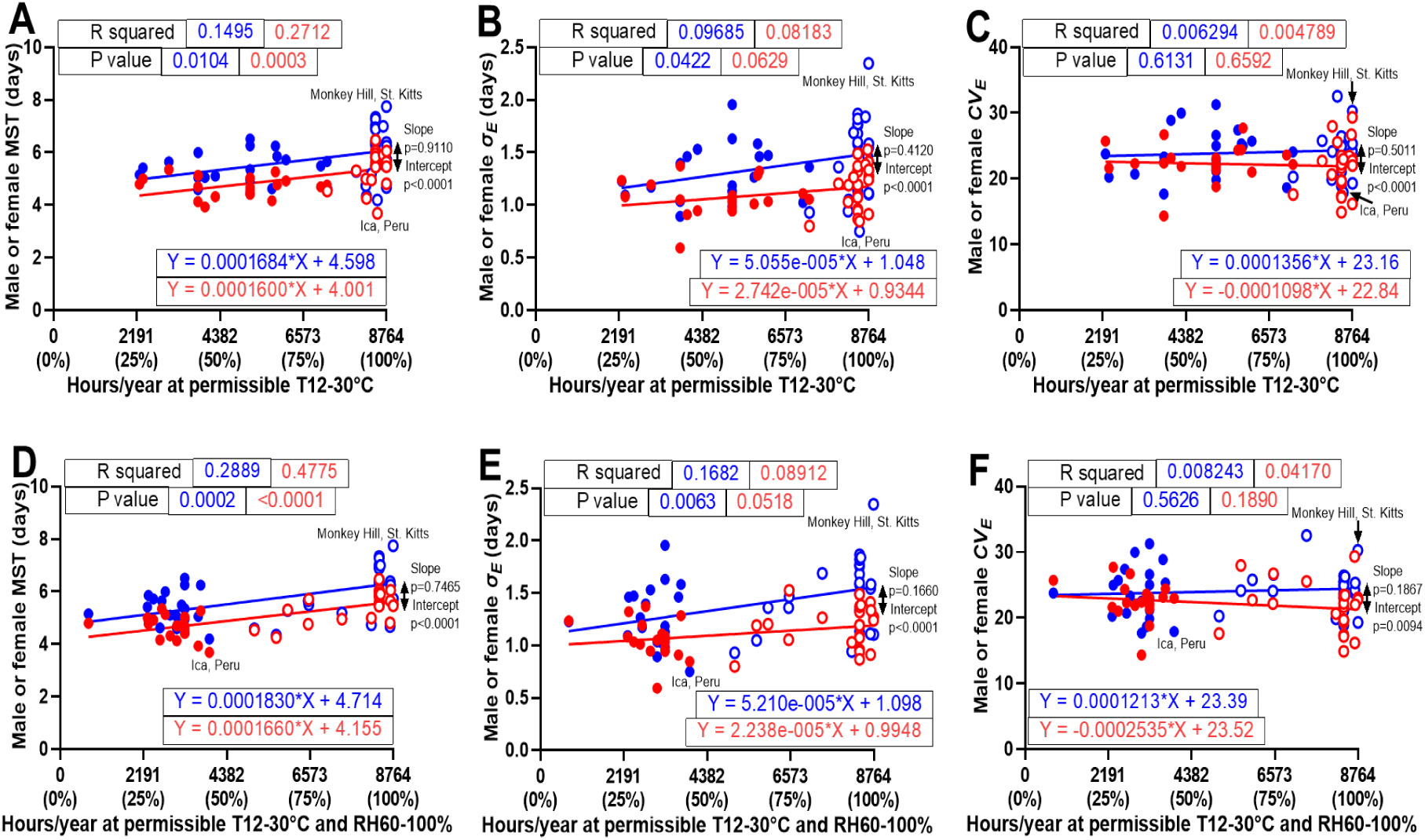
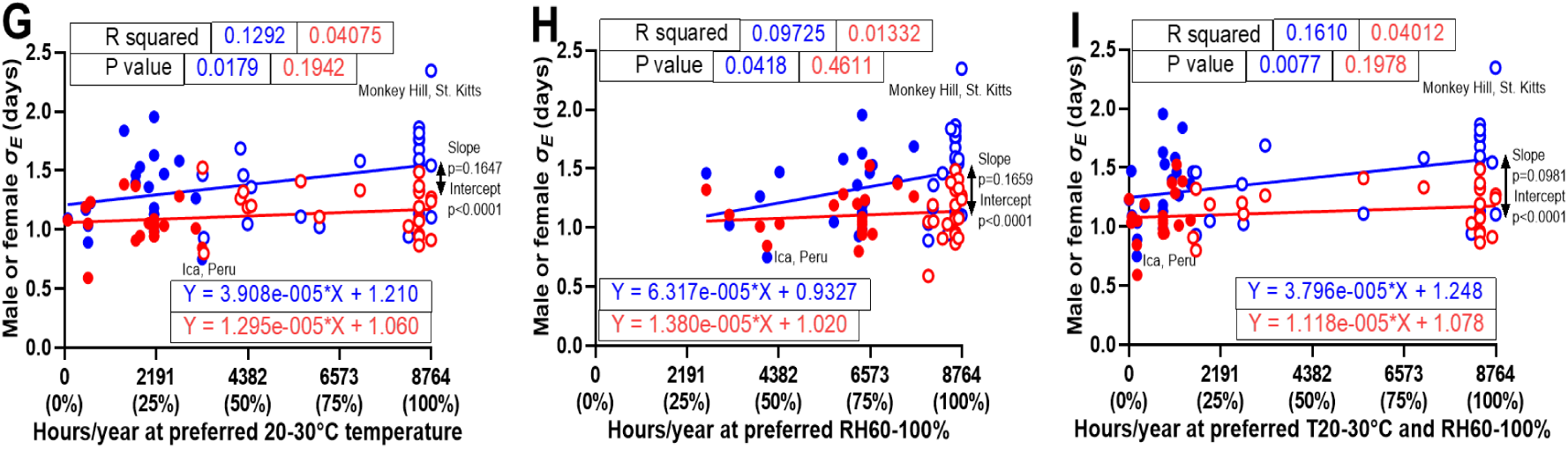
Simple linear regression analyses correlating Ma549 MST, *σ_E_*, and *CV_E_* values with hours of permissible temperature (12-30°C) with and without relative humidity (60-100%) or *σ_E_* values with preferred temperature (20-30°C), or preferred relative humidity (60-100%) or preferred temperature (20-30°C) coinciding with preferred relative humidity (60-100%) at collection sites for global fly populations. The analysis uses 52,584 hourly data points from 2005-2010 from all 28 sites for each fly line (S3A Table). Results are shown for permissible temperature with MST (**A**), *σ_E_* (**B**), and *CV_E_* (**C**), and for permissible temperature coinciding with relative humidity with MST (**D**), *σ_E_*(**E)**, and *CV_E_* (**F**), as well as *σ_E_*values with preferred temperature (**G**), or preferred relative humidity (**H**) or preferred temperature coinciding with preferred relative humidity (**I**) at collection sites for global fly populations, including R^2^ values, p-values, and regression equations with slopes. Male (female) regression lines are in blue (red). Male and female regression lines are blue and red, respectively. Aseasonal flies are depicted as circles, and seasonal flies as solid dots; this distinction applies to all subsequent figures. Males exhibited a positive correlation between *σ_E_* plasticity and exposure to permissible or preferred conditions (B, E, G-I).

**Figure 4—figure supplement 1.**
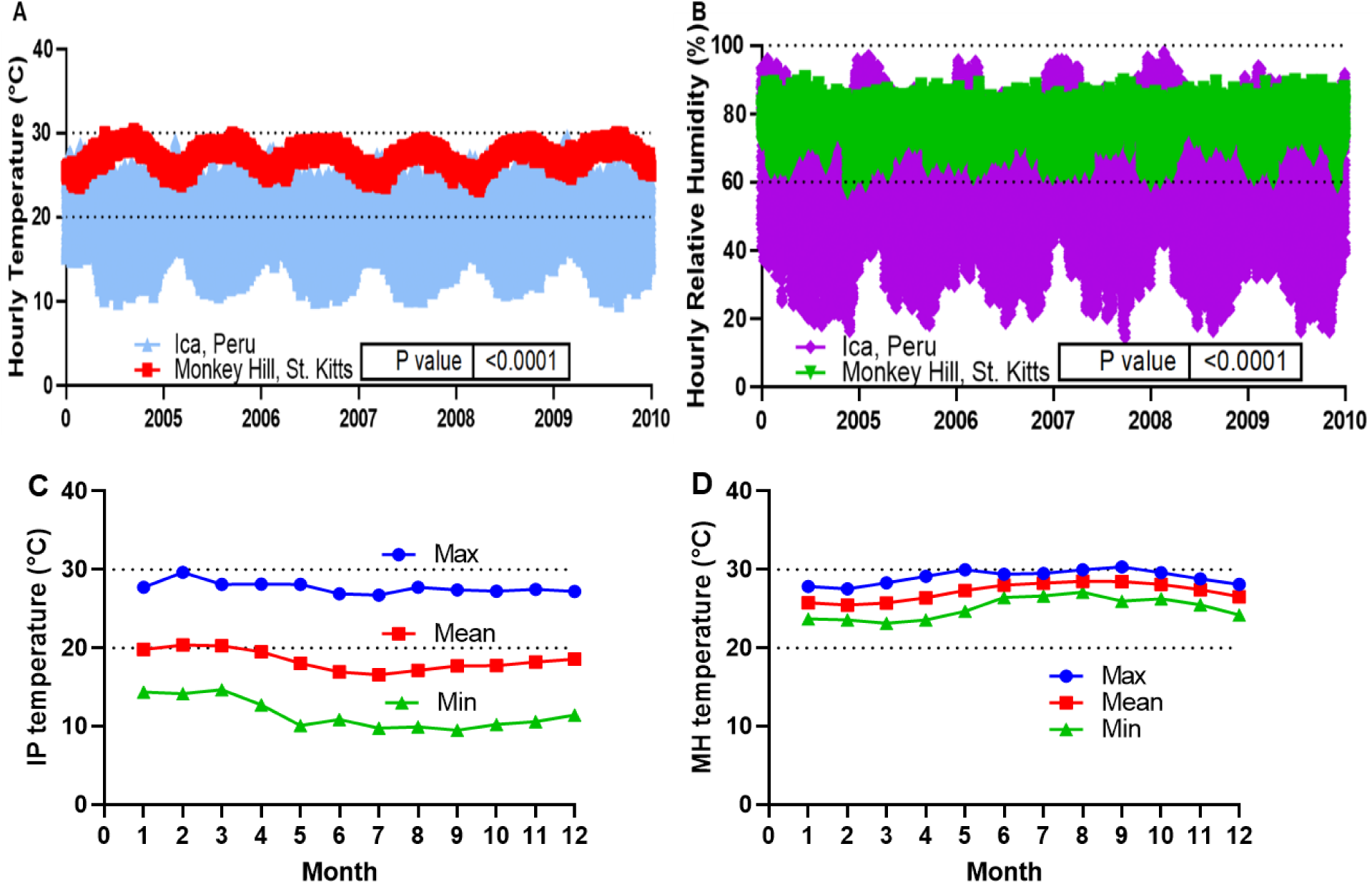
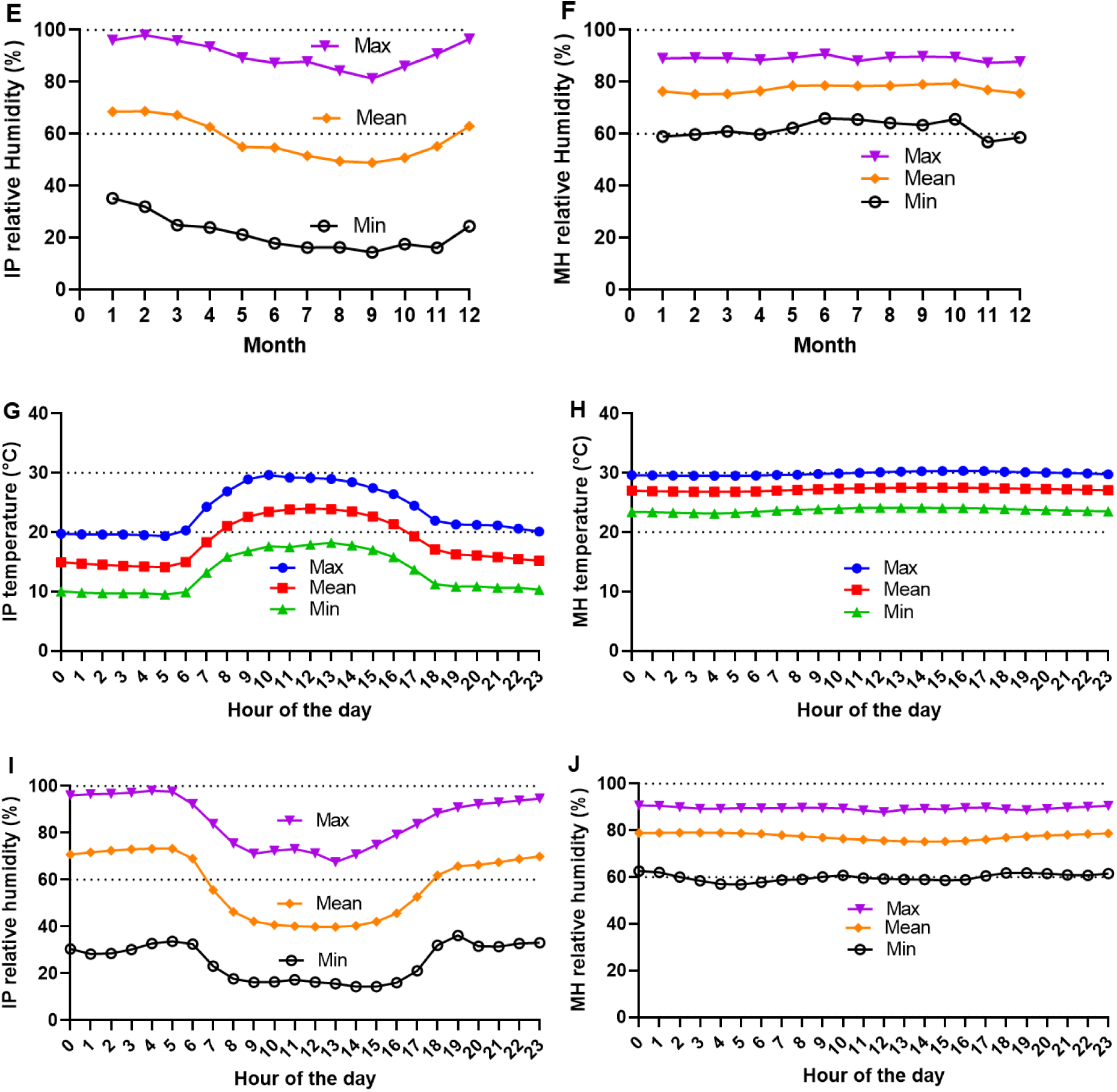
Hourly temperature **(A)** and relative humidity **(B)** data for Monkey Hills, St. Kitts (MH) and Ica, Peru (IP) are shown through Lomb-Scargle periodograms, capturing hourly fluctuations from January 01, 2005, to December 31, 2010 (n=52,584). Comparisons were made using Welch’s t-test with mean ± SEM and p-values indicated. Monthly temperature **(C, D)** and relative humidity **(E, F)** plots for IP **(C, E)** and MH **(D, F)** display maximum, mean, and minimum values across months from 2005-2010 (n=52,584 hourly data points), identifying seasonal trends and temperature-humidity interactions. Hourly temperatures (G, H) and relative humidity **(I, J)** plots for IP **(G, I)** and MH **(H, J)** illustrate daily patterns of maximum, mean, and minimum temperatures and relative humidity from 2005-2010 (n=52,584 hourly data points). These visuals help identify seasonal variations and trends in temperature and humidity interactions.

## Appendix 4—table 1 Effects of line, sex, mating status, and infection on sleep

ANOVA was used to validate the effects of genetic line, sex, mating status, and infection on nighttime (Figure 8—figure supplement 1A) and daytime (Figure 8—figure supplement 1B) sleep, as detailed in Table 1 (derived from the data in S5B Table). The analysis confirmed that infection status significantly affected daytime sleep (p < 0.0001) but not nighttime sleep (p = 0.5364), highlighting infection’s role in altering sleep at different times. Additionally, mating status had a much stronger impact on daytime sleep. A significant three-way interaction among line, infection, and sex (p ≤ 0.0020) was observed for both sleep periods, indicating that the influence of one factor on sleep can be significantly modified by the others.

**Appendix 4—table 1.** Effects of line, sex, mating status, and infection as well as their interactions on night and daytime sleep.

| Source | Nparm | D F | Nighttime sleep |  |  | Daytime sleep |  |  |
| --- | --- | --- | --- | --- | --- | --- | --- | --- |
|  |  |  | Sum of Squares | F Ratio | p value | Sum of Squares | F Ratio | p value |
| line | 2 | 2 | 4286291.9 | 874.5660 | <0.0001 | 1876451.8 | 219.7682 | <0.0001 |
| Infection | 1 | 1 | 937.4 | 0.3825 | 0.5364 | 3622098.1 | 848.4331 | <0.0001 |
| Mating | 1 | 1 | 17285.3 | 7.0537 | 0.0081 | 422367.3 | 98.9345 | <0.0001 |
| Sex | 1 | 1 | 1024007.7 | 417.8727 | <0.0001 | 17504.2 | 4.1001 | 0.0432 |
| line*Infection | 2 | 2 | 4138.2 | 0.8444 | 0.4302 | 34266.2 | 4.0132 | 0.0185 |
| line*Mating | 2 | 2 | 8143.3 | 1.6615 | 0.1906 | 2587.0 | 0.3030 | 0.7387 |
| line*Sex | 2 | 2 | 1519311.5 | 309.9971 | <0.0001 | 73329.8 | 8.5883 | 0.0002 |
| Infection*Mating | 1 | 1 | 2616.3 | 1.0676 | 0.3018 | 7005.3 | 1.6409 | 0.2006 |
| Infection*Sex | 1 | 1 | 132418.2 | 54.0366 | <0.0001 | 915803.3 | 214.5160 | <0.0001 |
| Mating*Sex | 1 | 1 | 694.8 | 0.2835 | 0.5945 | 22777.7 | 5.3354 | 0.0212 |
| line*Infection*Mating | 2 | 2 | 90.1 | 0.0184 | 0.9818 | 21402.3 | 2.5066 | 0.0822 |
| line*Infection*Sex | 2 | 2 | 30723.9 | 6.2688 | 0.0020 | 99378.2 | 11.6391 | <0.0001 |
| line*Mating*Sex | 2 | 2 | 1820.1 | 0.3714 | 0.6899 | 15843.5 | 1.8556 | 0.1571 |
| Infection*Mating*Sex | 1 | 1 | 1845.4 | 0.7531 | 0.3858 | 162.7 | 0.0381 | 0.8453 |
| line*Infection*Mating*Sex | 2 | 2 | 6.5 | 0.0013 | 0.9987 | 15603.2 | 1.8274 | 0.1615 |

(Source: This is a list of the tested factors and interactions. Nparm: Number of parameters associated with the effect. DF: degrees of freedom for each source of variation. Sum of Squares: sum of squares (SS) for each source of variation, along with the total from all sources. F Ratio: the mean square of the factor (lot) divided by the mean square of the error. A p-value less than 0.05 means that the factor or interaction has a statistically significant effect.)

**Figure 8—figure supplement 1.**
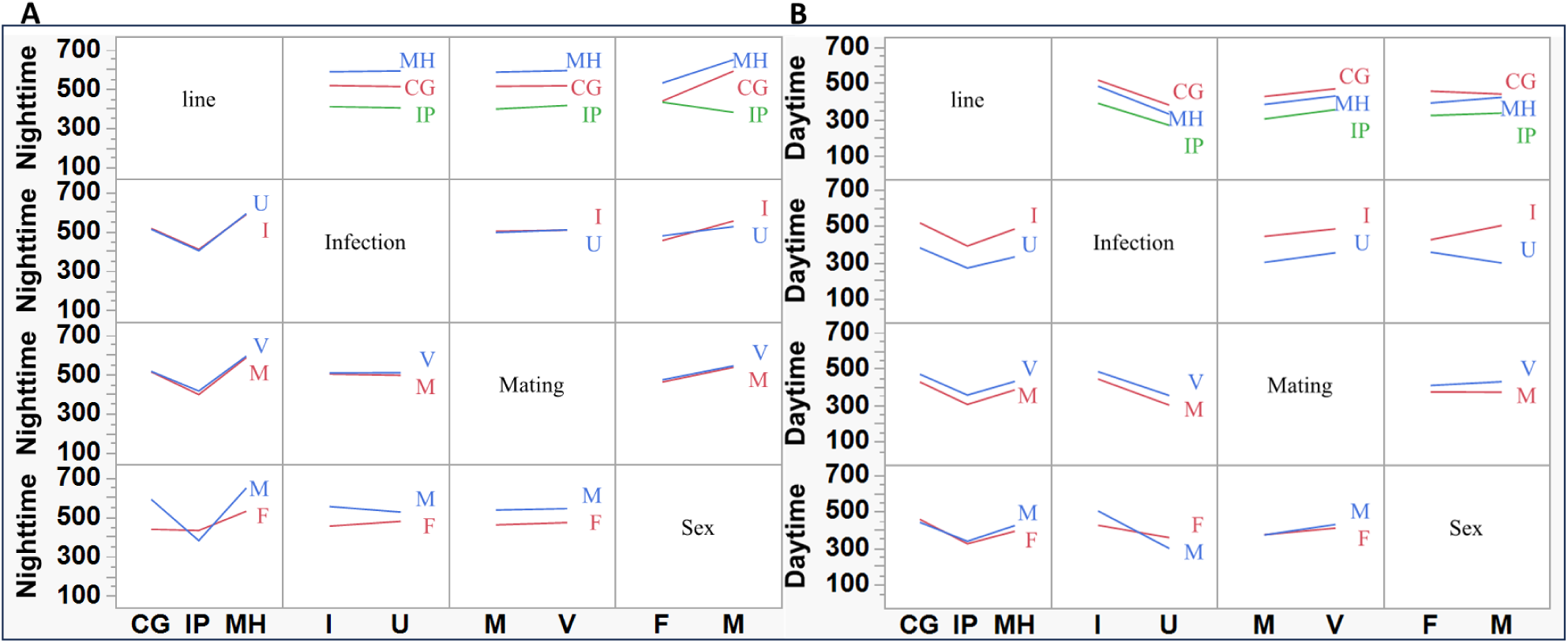
Interaction profiles of line, sex, mating status, and infection status on nighttime **(A)** and daytime **(B)** sleep (min/12h) across MH, CG, and IP females and males. These profiles were derived from the sleep data of 768 individual flies, considering three genetic lines (MH, CG, IP), both sexes (females and males), two mating statuses (mated and virgin), and two infection statuses (infected and uninfected). Abbreviations: sex (F = female, M = male), mating status (M = mated, V = virgin), and infection status (I = infected, U = uninfected).

